## Supplemental Data for "The *Legionella* collagen-like protein employs a unique binding mechanism for the recognition of host glycosaminoglycans"

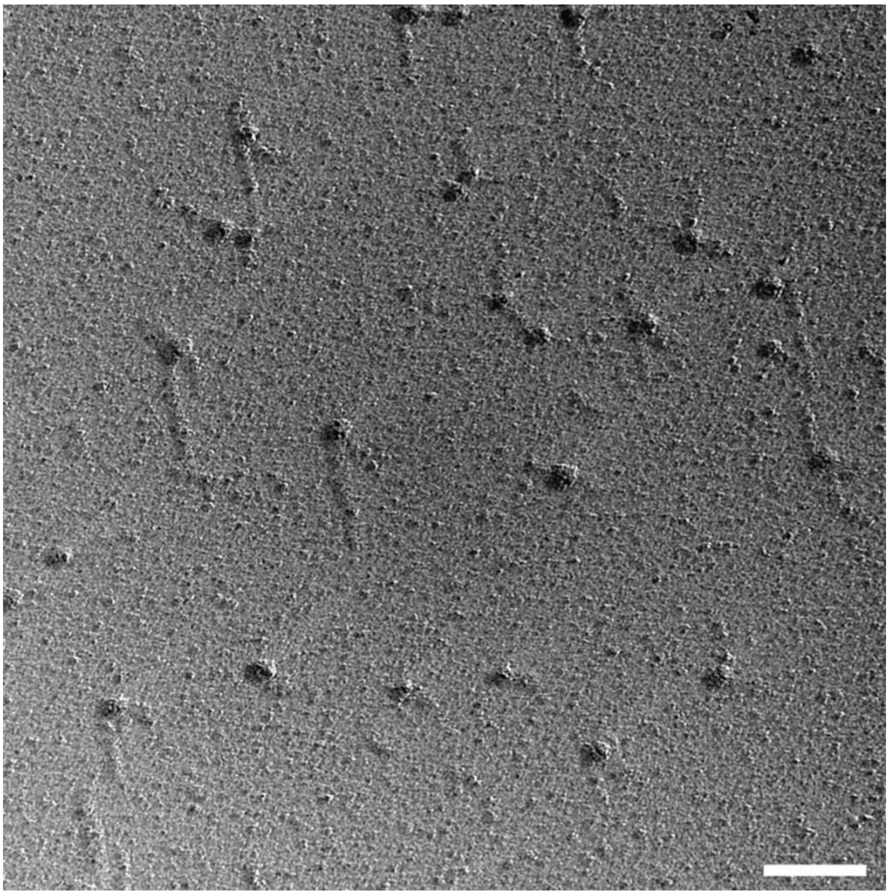

**Supplementary Fig. 1 | Rotary shadowing electron microscopy of Lcl.** Micrograph showing lollipop-shaped structures of Lcl trimers. The globular shapes correspond to trimeric C-terminal domains (CTD), while the stalks contain a trimeric collagen triple-helix (CLR). Some globular heads are missing the stalk region due to proteolysis. Variation in stalk lengths is due to differences in the platinum film thickness. The concentration of Lcl was 5 µg/ml. Scale bar: 50 nm.

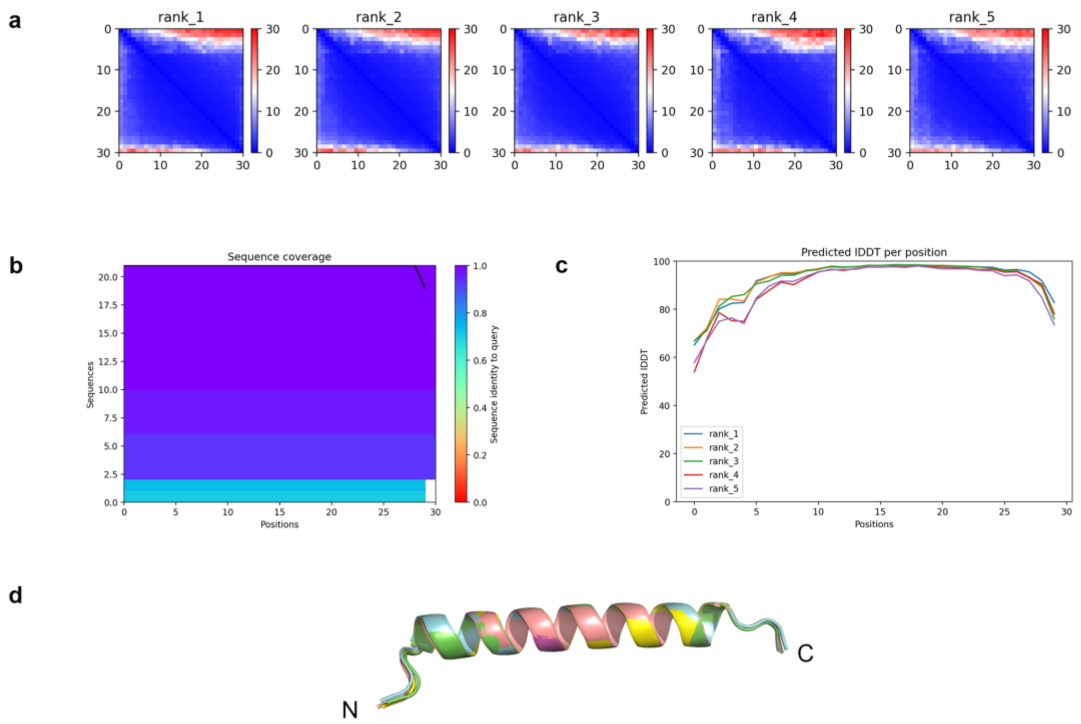


**Supplementary Fig. 2 | AlphaFold2 modelling of Lcl-N.** (a) Predicted Aligned Error (PAE) analysis of the five top ranking models. Low PAE values throughout the matrix indicates a well folded single domain. (b) Sequence coverage and identity used during modelling. (c) Local Distance Difference Test (lDDT) analysis against residue position for the five top ranking models. High lDDT values (<90) indicate a high accuracy in atomic positions for that residue. Lower values (<70) indicate low confidence atomic positioning, which is observed in dynamic regions. (d) Superposed top five ranked models with the termini highlighted (N/C: N-/C-terminus).

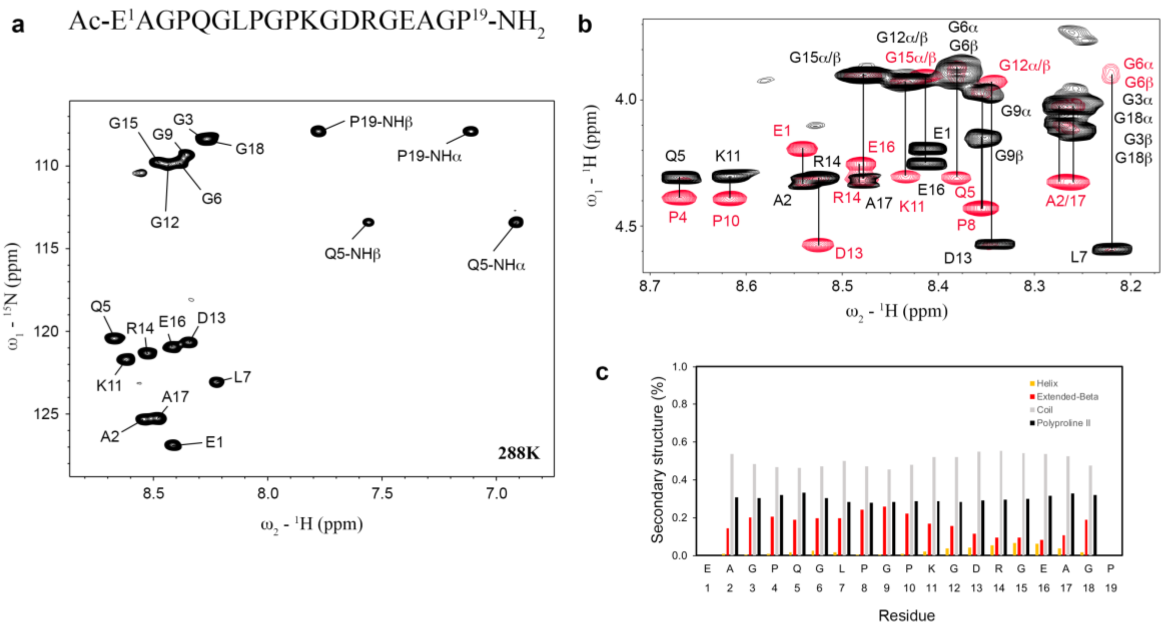


**Supplementary Fig. 3 | Solution NMR spectroscopy analyses of unlabelled CLR peptide recorded at 15°C. a,** $^{1}H$–$^{15}N$ SOFAST-HMQC spectrum of the CLR peptide with amide resonance assignments shown. Peptide sequence is shown above. **b,** $^{1}H$–$^{1}H$ TOCSY (black) and $^{1}H$–$^{1}H$ ROESY (red) spectra expanded on the NH-$H_\alpha$ region. A typical pattern of polyproline II structure is seen with strong NOE correlation observed between NH (i) and $H_\alpha$ (i-1). **c,** Secondary structure propensity of monomeric CLR peptide derived from backbone chemical shifts ($C_\alpha$, $C_\beta$, $H_\alpha$, N, NH) calculated using δ2D[1].

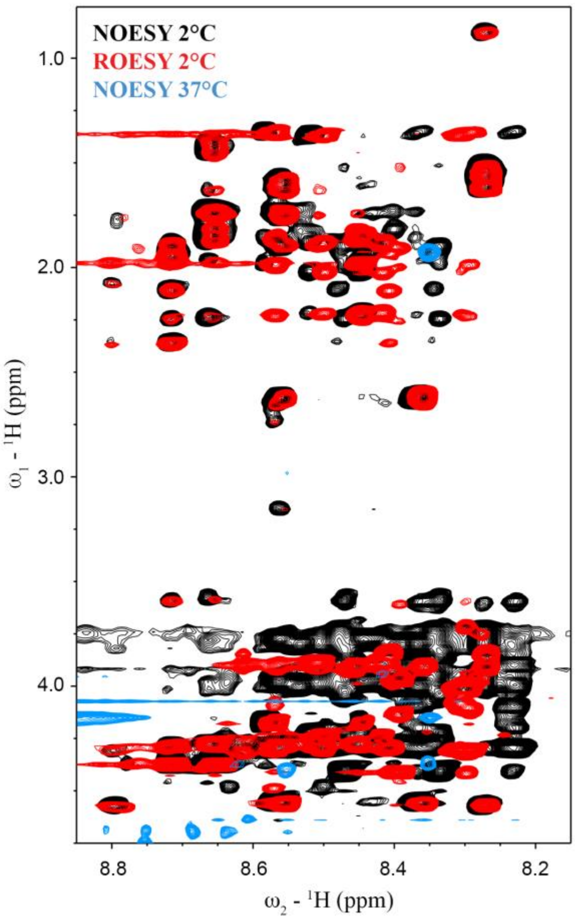


**Supplementary Fig. 4 | Solution NMR spectroscopy analyses of $^{13}C^{15}N$ glycine labelled CLR peptide.** $^{1}H$–$^{1}H$ NOESY spectra (mixing time 240 ms) recorded at 2°C (black) and 37°C (blue), and $^{1}H$–$^{1}H$ ROESY (mixing time 200 ms) spectrum recorded at 2°C (red) expanded on the amide region. Significant differences are observed between the NOESY and ROESY spectra at 2°C and significant spectral broadening is observed in the NOESY spectrum at 37°C. This indicates that the peptide is in equilibrium between monomeric and trimeric states.

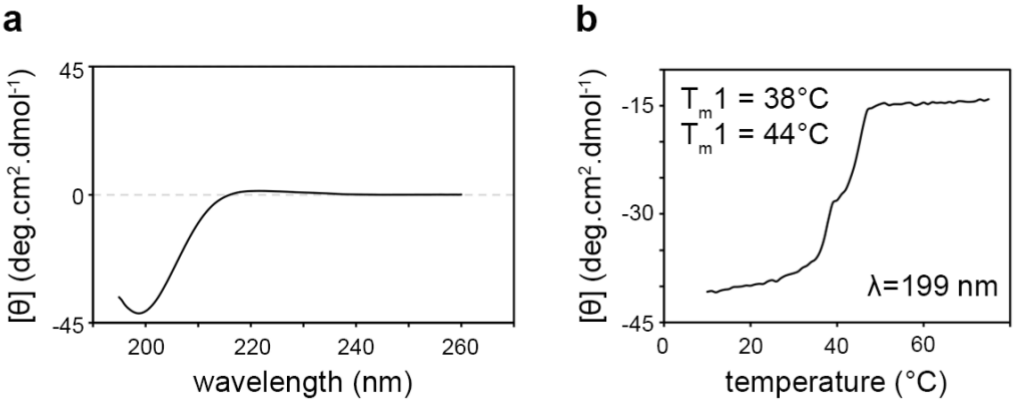


**Supplementary Fig. 5 | Circular dichroism (CD) spectra of Lcl. a,** CD spectra of Lcl recorded at 10°C with negative peak at 199 nm and maximum peak at 222 nm. **b,** Thermal denaturation of Lcl recorded between 10°C and 75°C at 199 nm. Two melting temperatures were determined at 38°C and 44°C.

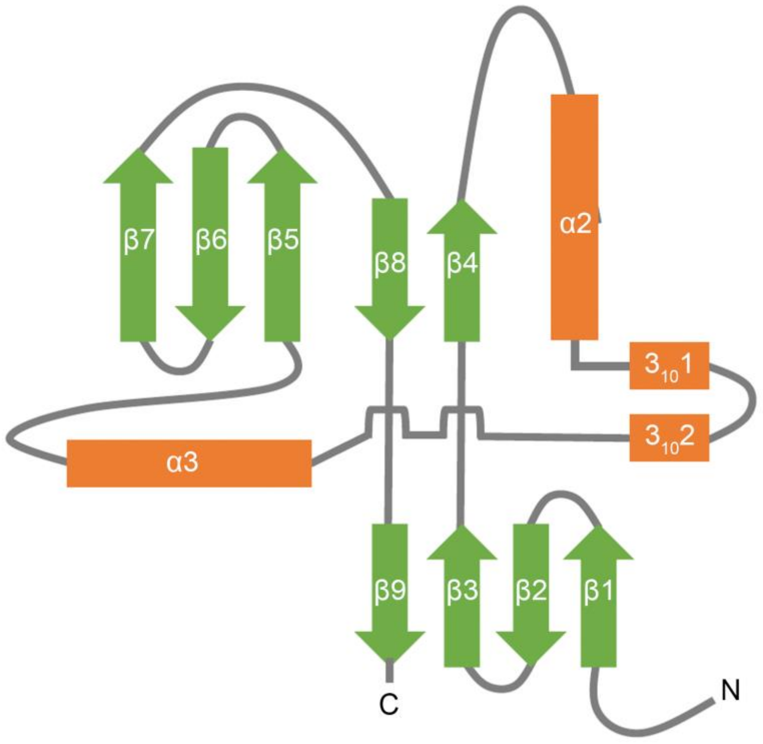


**Supplementary Fig. 6 | Topology diagram for Lcl-CTD.** Schematic representation of an individual Lcl-CTD domain. α-helix and $3_{10}$-helix secondary structure are coloured orange, β-sheet secondary structure is coloured green, and loops are grey. Helices are labelled 1-2 and β-sheets are labelled 1-9.

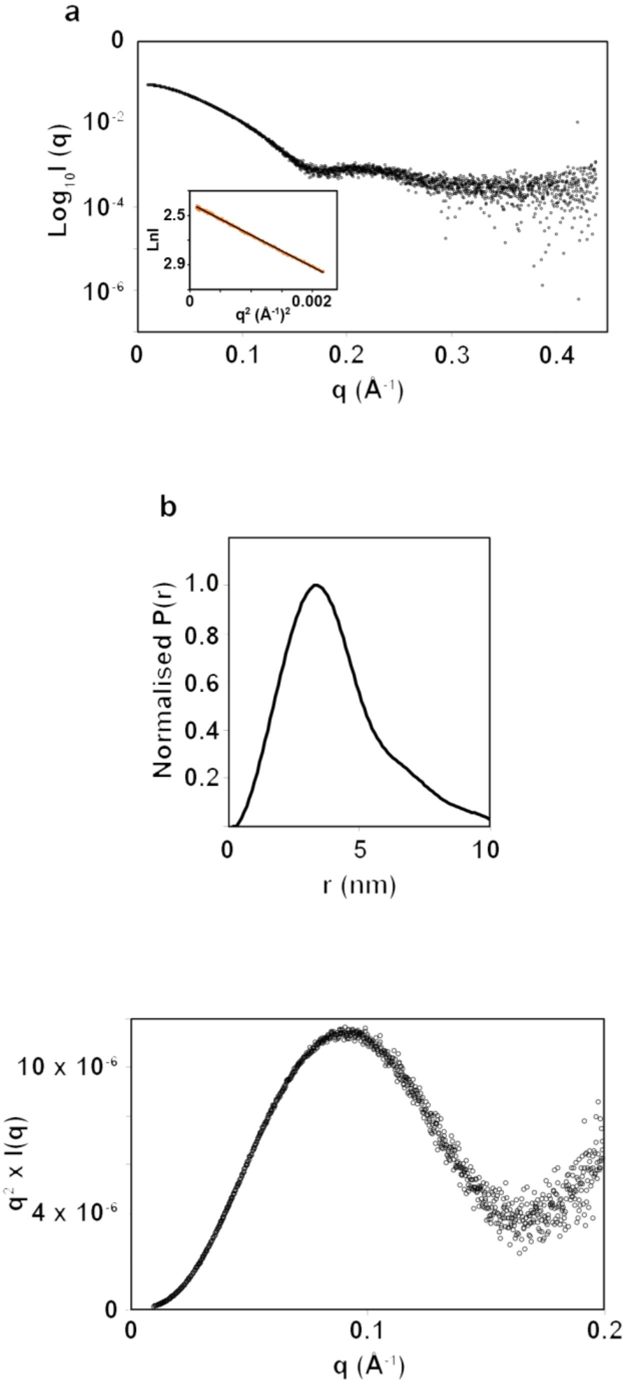


**Supplementary Fig. 7 | SAXS analysis of Lcl-CTD. a,** Experimental scattering curve of Lcl-CTD (black open circles). Inset: Guinier region (orange open circles) and linear regression (black line) for $R_g$ evaluation. **b,** Shape distribution [P(r)] function derived from SAXS analysis for Lcl-CTD. **c,** Kratky, plot indicates that Lcl-CTD has dynamic properties in solution.

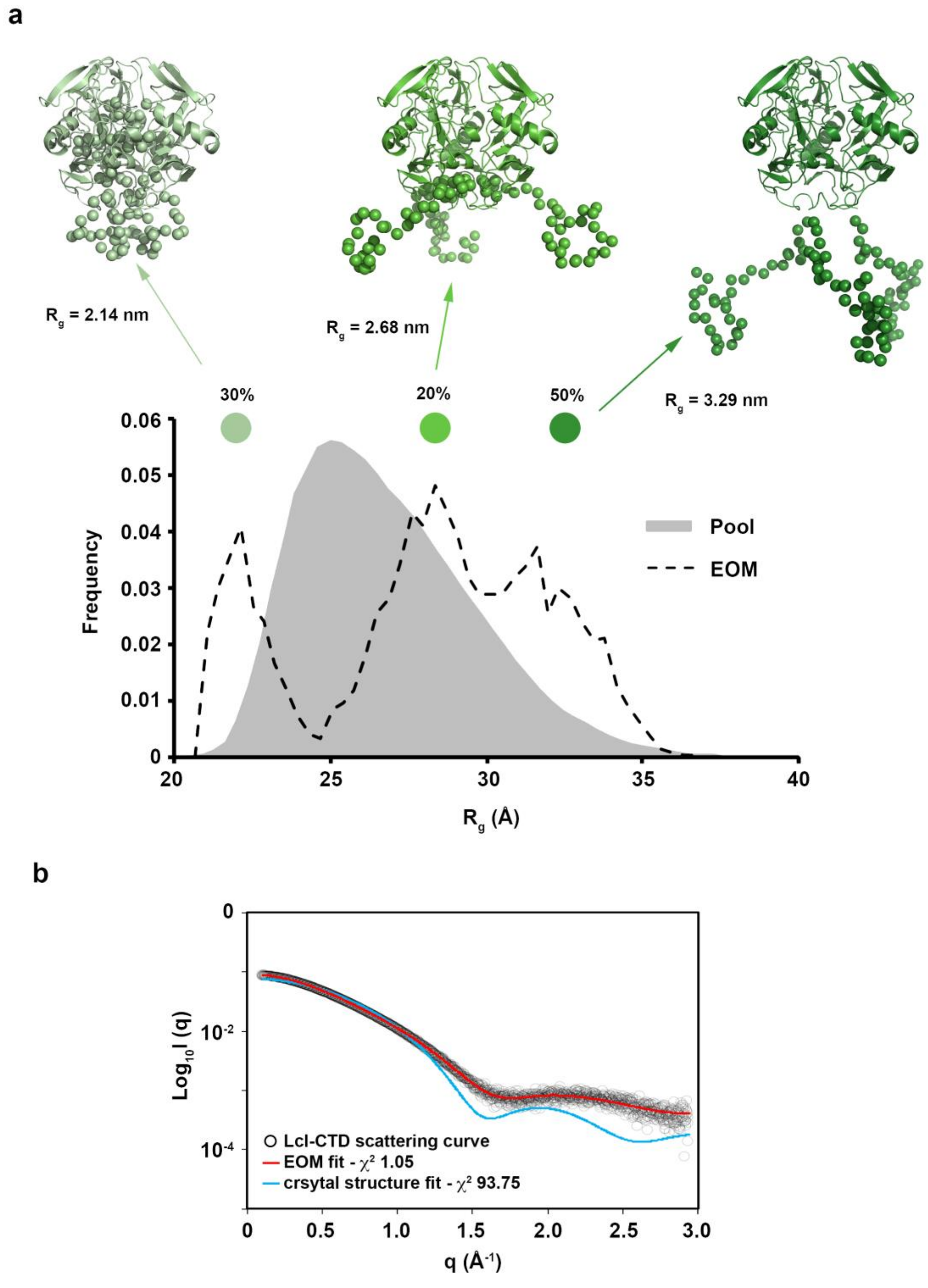


**Supplementary Fig. 8 | SAXS derived model of Lcl-CTD in solution. a,** EOM yielded three distinct populations: extended N-terminus (50%), partially extended N-terminus (20%) and compact N-terminus (30%). Lcl-CTD models corresponding to the centre of each population are shown. **b,** EOM (red line) and crystal structure (blue) fit to the Lcl-CTD SAXS data (black open circles) with $\chi^2$ of 1.05 and 93.75, respectively.

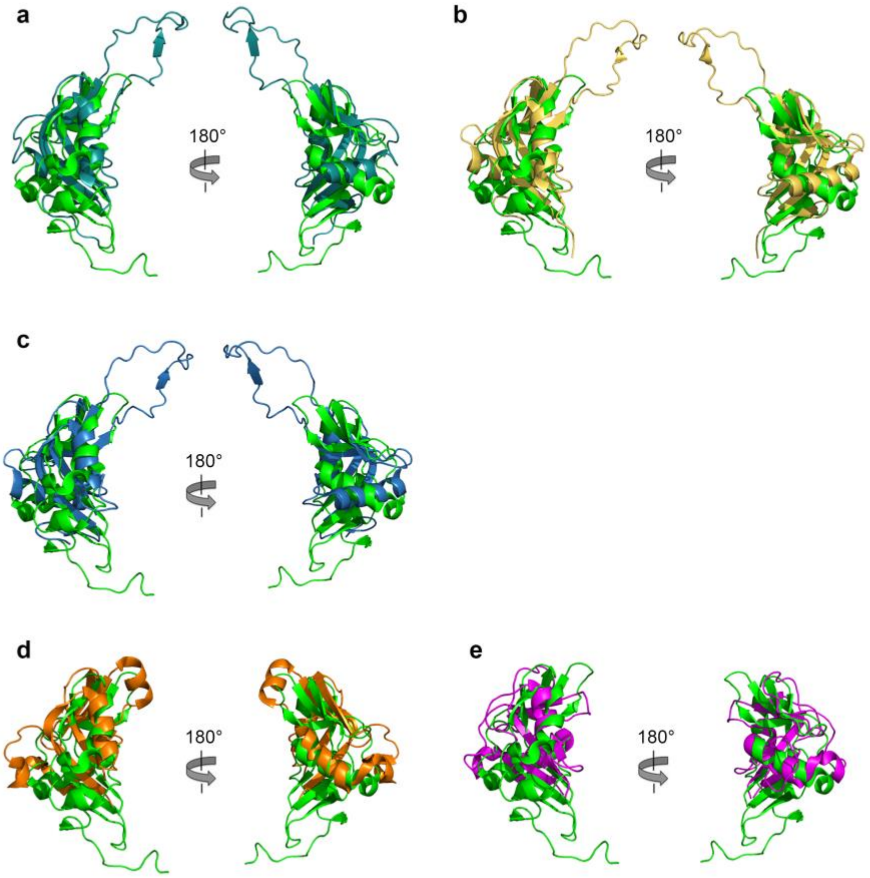


**Supplementary Fig. 9 | Structural comparison of the Lcl-CTD monomer with other C-type lectin-like domains.** Homologous structures identified using the Dali server with Z-scores >8.0 are shown[2]. **a,** Cartoon representation of Lcl-CTD (green) superimposed on the platelet glycoprotein Ib-α binding protein mucrocetin from the venom of *Trimeresurus mucrosquamatus* (teal; Protein Data Bank (PDB) ID code 1v4l[3]; chain B residues 201–325; Z-score 9.7; root mean squared deviated (rmsd) 2.7 Å). **b,** Cartoon representation of Lcl-CTD (green) superimposed on the integrin binding protein EMS16 from the venom of *Echis multisquamatus* (yellow; PDB ID code 1v7p[4]; chain A residues 1–127; Z-score 9.1; rmsd 2.7 Å). **c,** Cartoon representation of Lcl-CTD (green) superimposed on the integrin binding protein rhodocetin from the venom of *Calloselasma rhodostoma* (blue; PDB ID code 6nd8; chain B residues 1–122; Z-score 8.5; rmsd 2.8 Å). **d,** Cartoon representation of Lcl-CTD (green) superimposed on the adhesin protein intimin from enteropathogenic *Escherichia coli* (orange; PDB ID code 1f00[5]; chain I residues 840–939; Z-score 8.3; rmsd 3.5 Å). **e,** Cartoon representation of Lcl-CTD (green) superimposed on the integrin binding protein invasin from *Yersinia pseudotuberculosis* (purple; PDB ID code 1cwv[6]; chain A residues 887–986; Z-score 8.1; rmsd 2.9 Å).

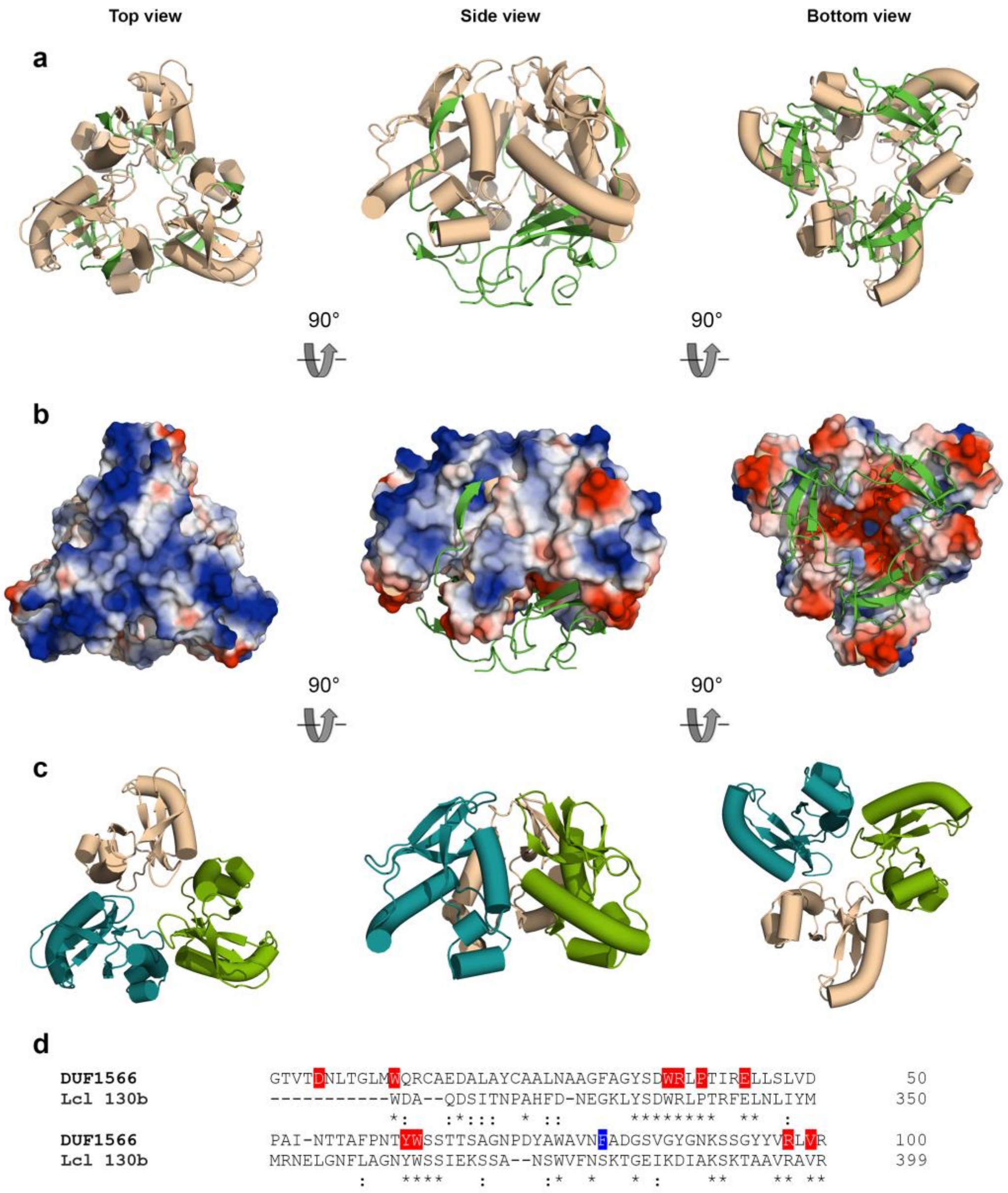


**Supplementary Fig. 10 | DUF1566/pfam07603 motif within Lcl-CTD. a,** Cartoon representation of trimeric Lcl-CTD with the DUF1566 regions coloured wheat, and the non-DUF1566 S1-S4 strands of Lcl-CTD coloured green. **b,** Electrostatic surface potential representation of the DUF1566 region within the Lcl-CTD trimer, with the Lcl-CTD S1-S4 strands shown as green cartoon. **c,** Cartoon representation of the isolated DUF1566 trimer from Lcl-CTD. **d,** Sequence alignment of the consensus DUF1566 motif and the corresponding region within *L. pneumophila* 130b Lcl. Identical residues are highlighted with an asterisk (*), and residues that are similar are highlighted with a colon (:). Completely conserved positions in the DUF1566 sequence that are also identical in Lcl are coloured red, and those that are not identical are coloured blue.

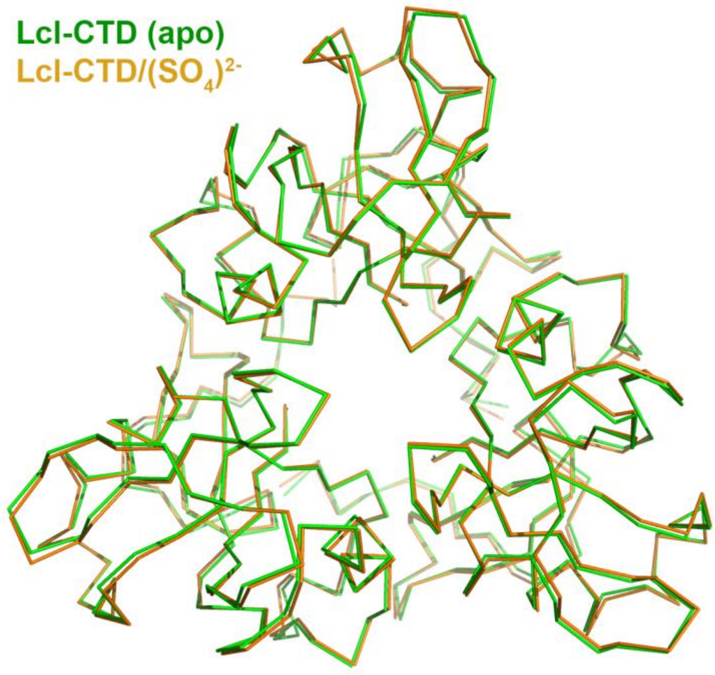


**Supplementary Fig. 11 | Structural comparison of the Lcl-CTD trimer is the presence and absence of sulphate ions.** Lcl-CTD trimer backbone structures are shown as ribbons in green (no sulphate) and orange (bound to sulphate ions).

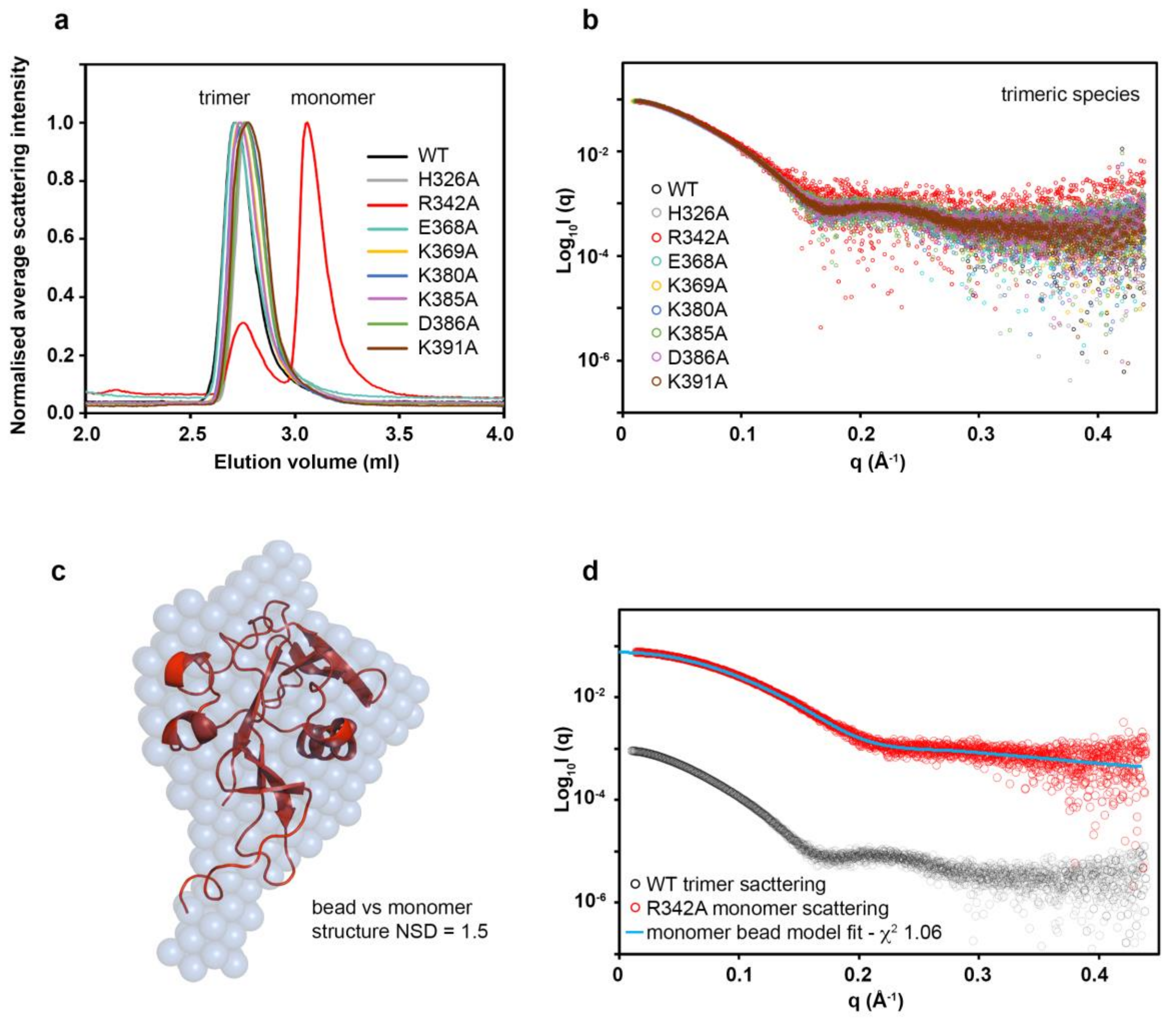


**Supplementary Fig. 12 | SAXS analysis of Lcl-CTD mutants. a,** Size-exclusion chromatography coupled with SAXS profile of wild-type and engineered Lcl-CTD. **b,** Experimental scattering curves of trimeric forms of wild-type and engineered Lcl-CTD. **c,** DAMMIF bead model of monomeric Lcl-CTD R342A superimposed with chain A from Lcl-CTD, with normalized spatial discrepancy (NSD) score of 1.5. **d,** Monomer Lcl-CTD crystal structure fit to monomeric Lcl-CTD R342A SAXS data (red open circles) with $\chi^2$ of 1.06. Experimental scattering curve of trimeric wild-type Lcl-CTD (black open circles) is shown for comparison.

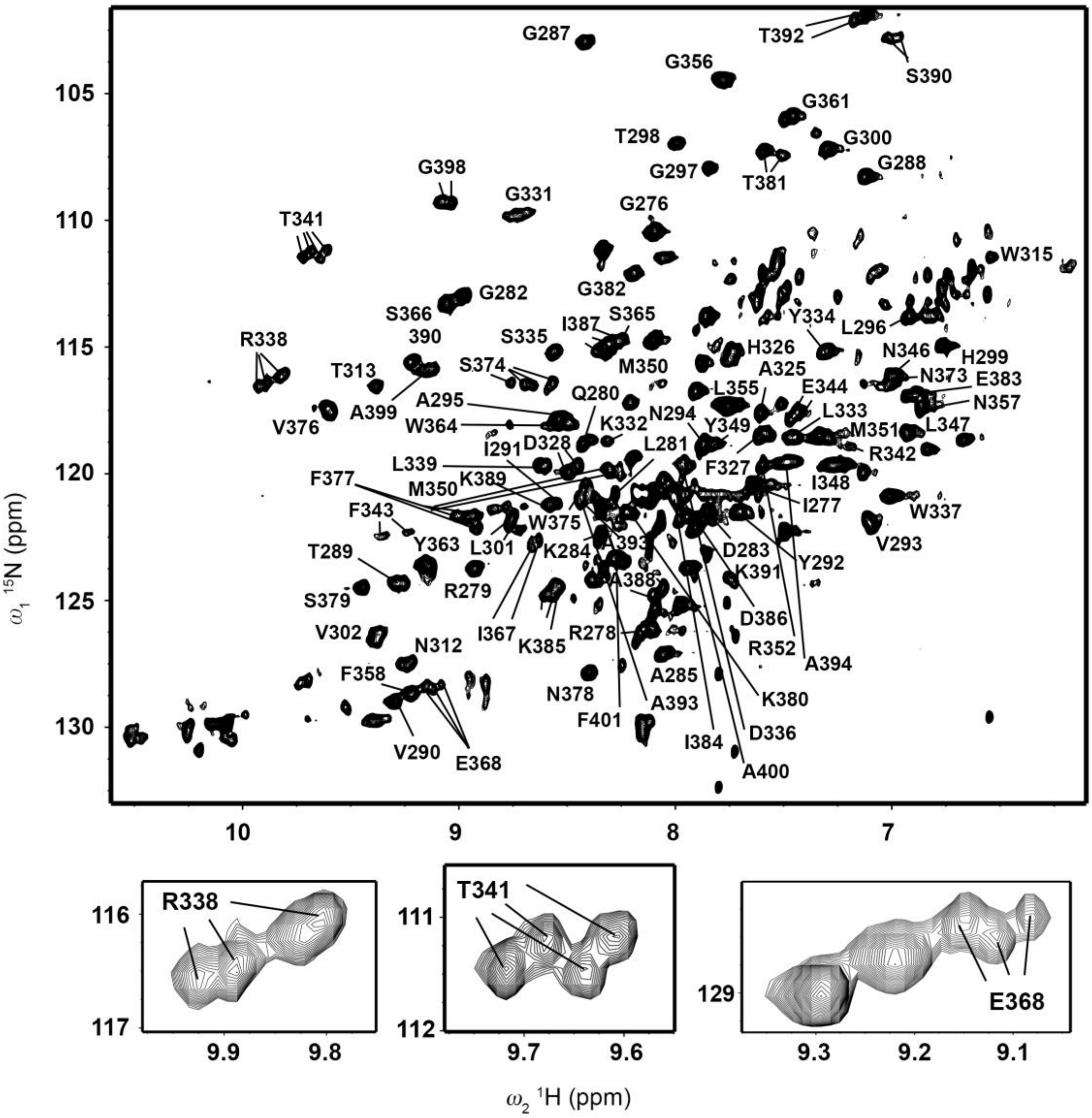


**Supplementary Fig. 13 | Solution NMR spectroscopy of the Lcl C-terminal domain.** $^{1}$H–$^{15}$N HSQC TROSY spectrum with backbone amide resonance assignments shown. Boxed out areas are examples of residues that display conformation exchange.

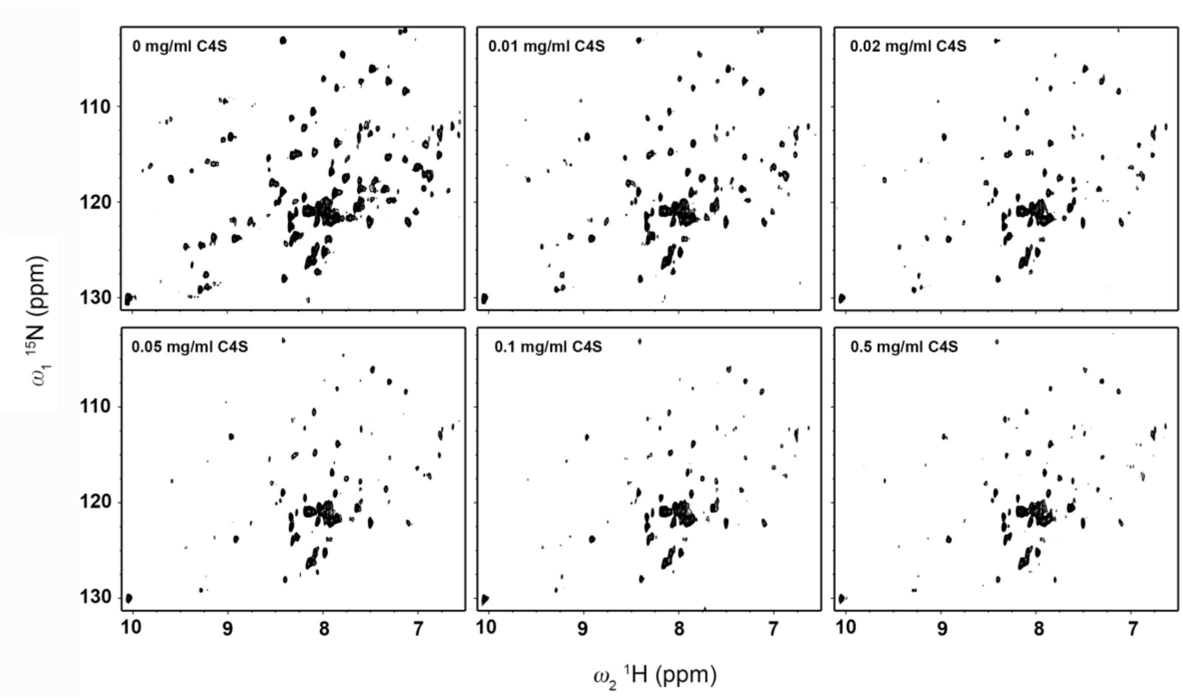


**Supplementary Fig. 14 | NMR titration of C4S against Lcl-CTD.** $^{1}H$–$^{15}N$ HSQC TROSY spectra of $^{2}H^{15}N$ labelled Lcl-CTD incubated with increasing concentrations of C4S extracted from bovine trachea.

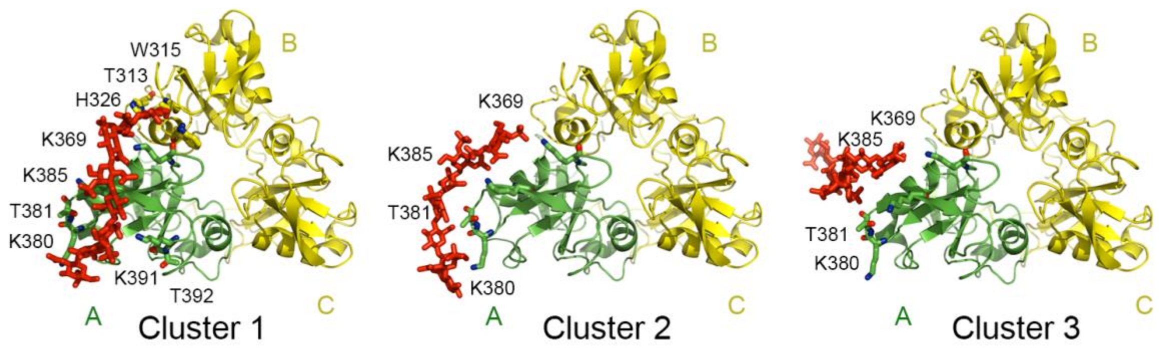


**Supplementary Fig. 15 | Docking of C4S dp8 against monomeric Lcl-CTD.** Top three HADDOCK clusters are shown. C4S is shown as sticks and coloured red, while monomeric Lcl-CTD is shown as a green cartoon. Docking was carried out on monomeric Lcl-CTD, but here it is shown superimposed onto the crystal trimer (yellow). The trimeric structure shown in Cluster 1 is equivalent to the HM model used for subsequent MD simulations.

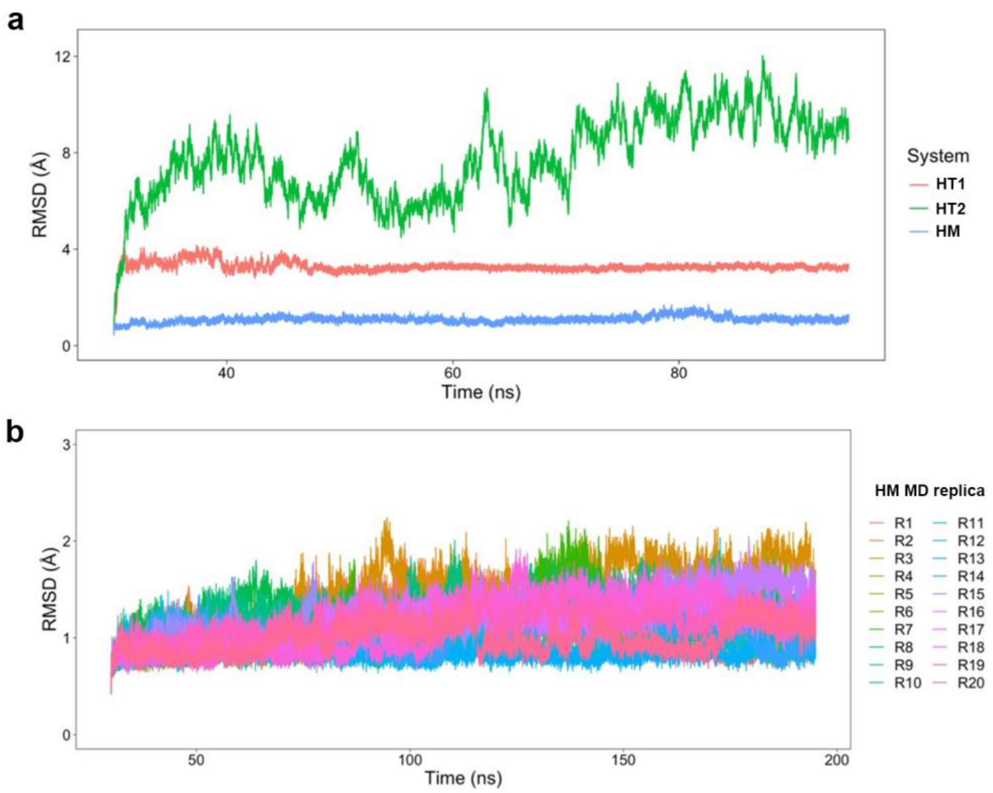


**Supplementary Fig. 16 | Stability of Lcl-CTD/C4S during Molecular Dynamics simulations. a,** Time evolutions of the RMSD of $C_\alpha$ atoms from the starting structure before energy minimisation for Lcl-CTD in complex with C4S. The HT1 and HT2 plots represent MD simulations starting from different HADDOCK models where one molecule of C4S dp8 was docked against a trimer of Lcl-CTD (3:1 Lcl-CTD:C4S). The HM plot represents a simulation starting from the HADDOCK model used for subsequent analysis. During HADDOCK one molecule of C4S dp8 was docked against a monomer (chain A) of Lcl-CTD (1:1 Lcl-CTD:C4S) and then prior to running the MD simulations it was reconstituted as a trimer (3:1 Lcl-CTD:C4S). **b,** The RMSD values for all the replicas starting from the HM model. For **a** and **b**, RMSD values were calculated after best-fit superimposition of each frame to the reference structure and the highly flexible N-terminal residues (271-277) were not included in the calculation. The initial equilibration stages with positional restraints were omitted from the plot.

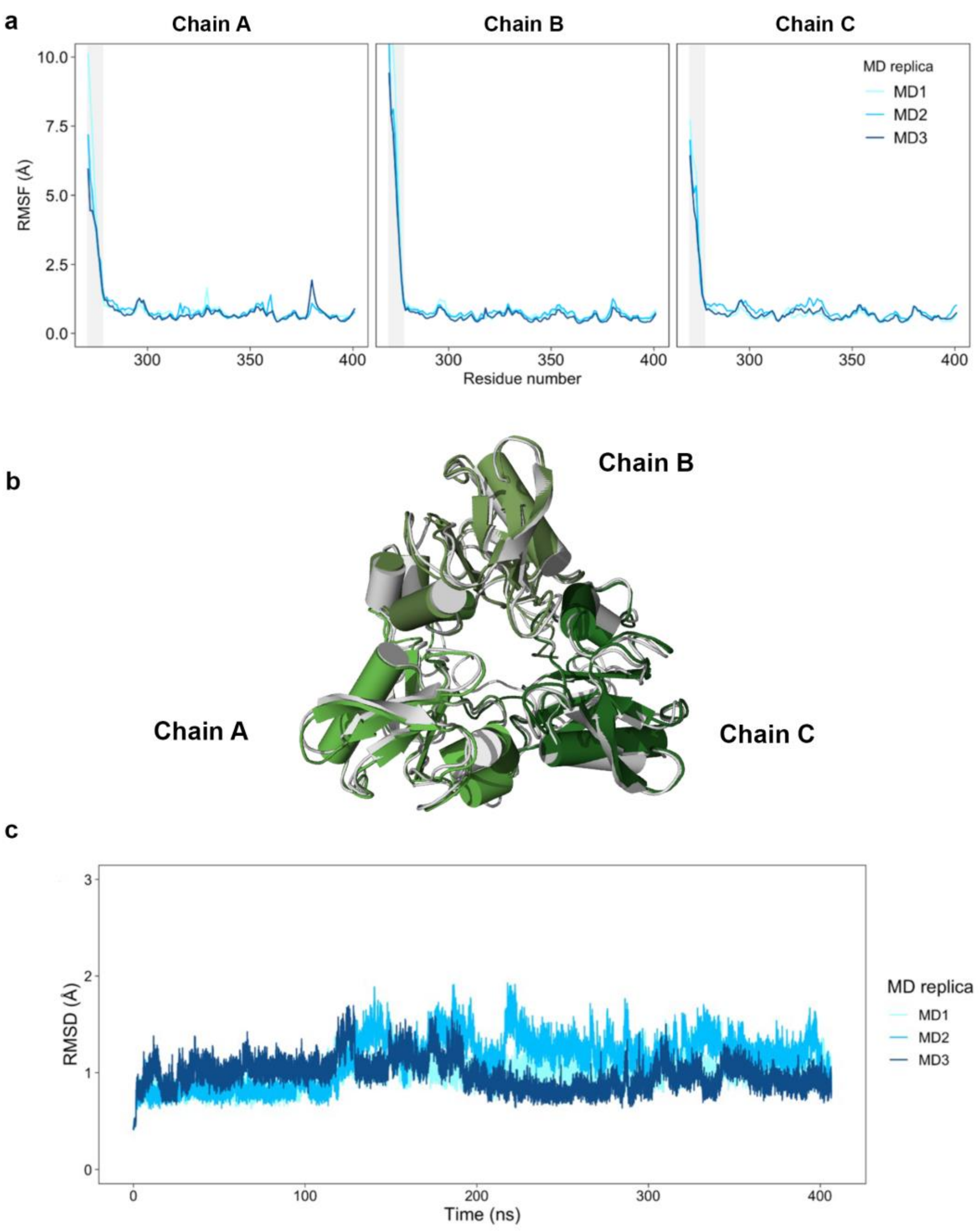


**Supplementary Fig. 17 | Molecular Dynamics simulations of Lcl-CTD. a,** RMSF profiles for each monomer and each replica of the Lcl-CTD simulations. The grey area highlights the position of the first 7 residues (271 to 277) for each monomer. RMSF values are calculated considering only $C_\alpha$ atoms and on frames saved every 1 ps. **b,** Representative structure of the most populated cluster (white, population=79.7%) from the MD simulations superimposed to the initial crystal structure (green hues). **c,** Time evolution of the RMSD of $C_\alpha$ atoms from the starting structure before energy minimisation for Lcl-CTD alone.

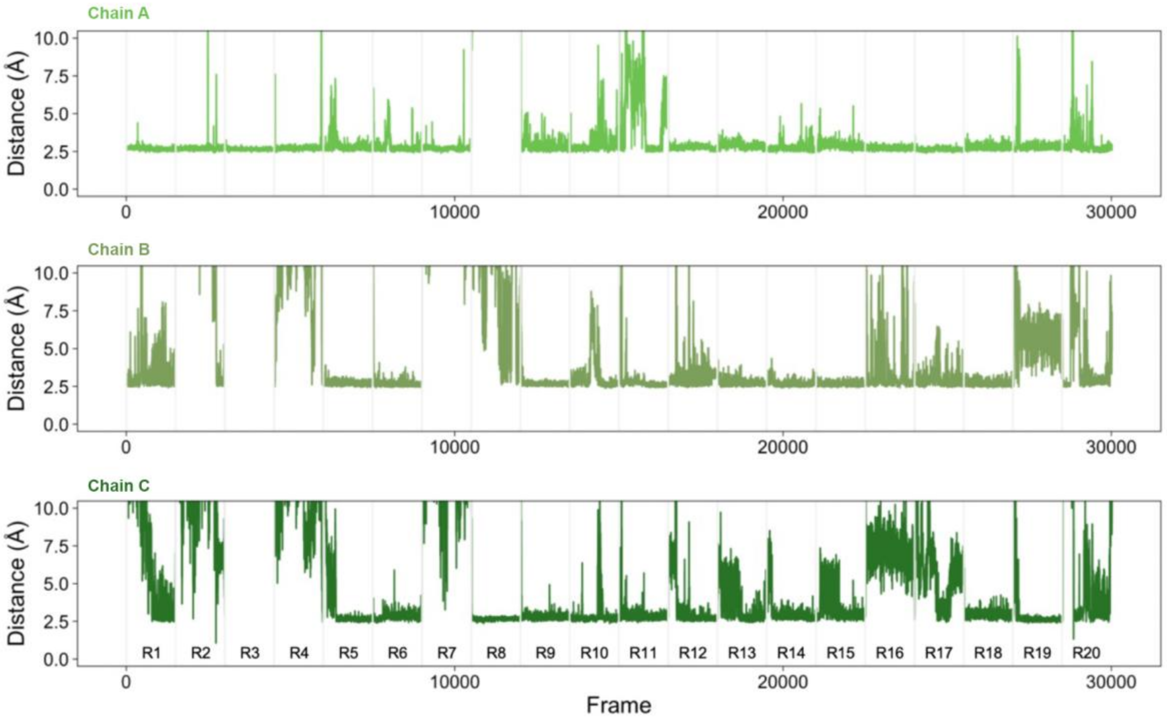


**Supplementary Fig. 18 | C4S binding to Lcl-CTD.** Time evolution of the distance between C4S and each chain of the Lcl-CTD trimer during the 20 MD replicas (production only). A grey vertical line separates different replicas. The distance was calculated as the minimum distance between all possible pairs of non-hydrogen atoms from C4S and Lcl-CTD. Analysis of the distances shows that C4S can be in contact with a single chain (36.2% of the frames from all replicas), 2 chains (34.8%) or across the whole trimer (27.7%).

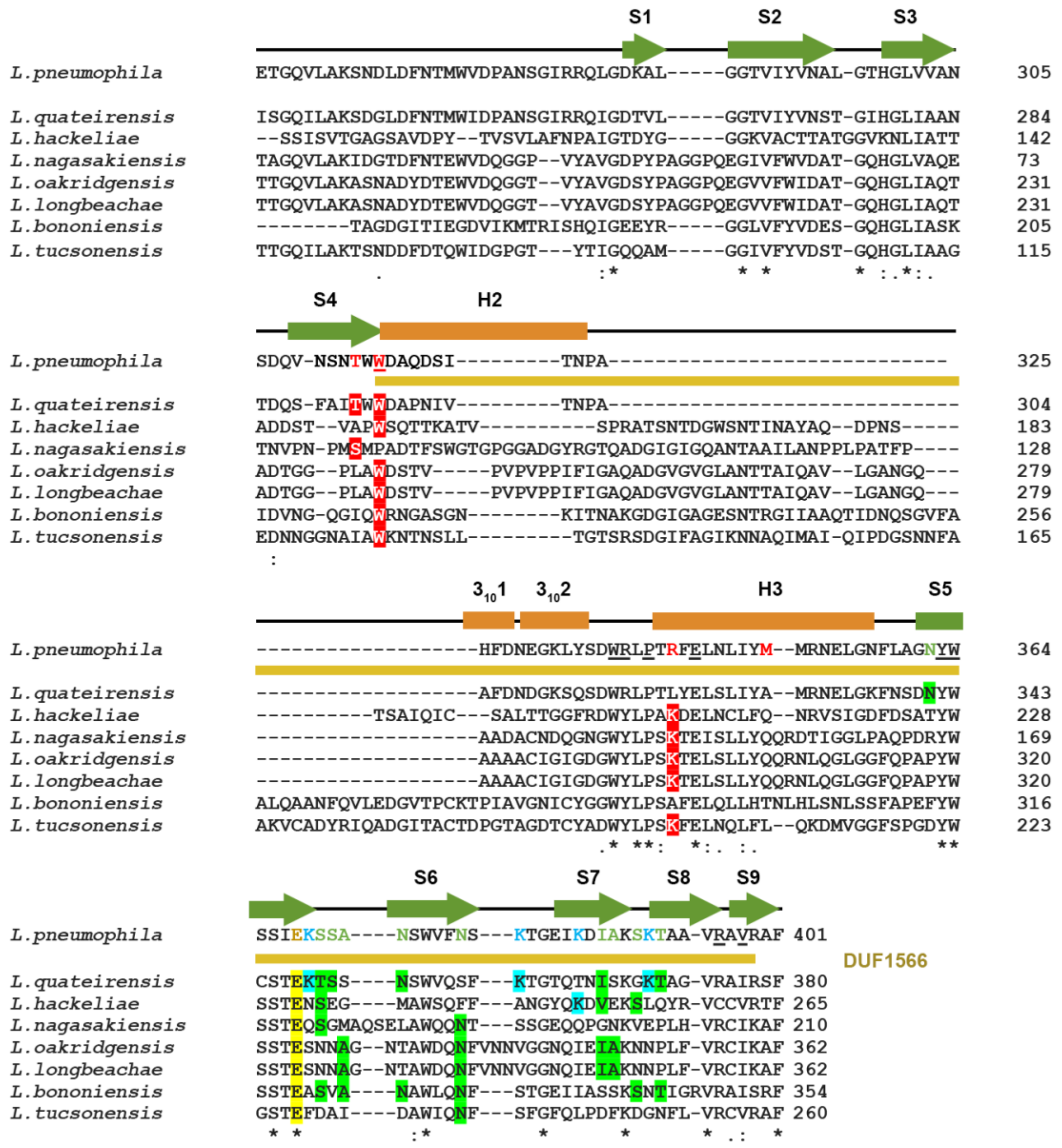

**Supplementary Fig. 19 | Sequence alignment of Lcl across the *Legionella* genus.** Amino acid positions with 100% identical, >50% identical and similar residues are indicated by asterisk (*), colon (:), and period (.), respectively. Residues in *L. pneumophila* 130b Lcl-CTD that influence GAG binding are coloured red (identified by NMR), green (identified by MD), yellow (internal glutamate; identified by ELISA) and blue (surface lysines; identified by ELISA). Identical residues in other species are shaded in the same colour. Secondary structure elements and the position of DUF1566 are shown for *L. pneumophila* 130b Lcl-CTD, with related residues in DUF1566 that are highly conserved, underlined in the 130b sequence.

**Supplementary Table 1 | Domain boundaries of Lcl from *L. pneumophila* 130b strain**

The predicted N-terminal amphipathic helix is underlined with residues predicted to form the hydrophobic face in bold[7].

| | |
|---|---|
| Flexible N-terminus<br>(residues 1 to 30) | KSN<u>PASQAY**V**DGK**V**SE**L**KNE**L**TNK**I**N</u>SIPS |
| CLR region<br>(residues 31 to 251) | GPQGPRGDKGEAGPKGDRGEAGPQGLPGPKGDRGEAGPQGLPGPKGDRGEAGPQGLPGPKGDRG<br>EAGPQGLPGPKGDRGEAGPQGLPGPQGLPGPKGDKGEAGPQGLPGPKGDKGEAGPQGLPGPKGD<br>KGEAGAVGPQGMPGPKGDKGEAGPQGLPGPKGDRGEAGPQGLPGPKGDRGEAGPQGLPGPKGDK<br>GETGAVGPQGMPGPKGEAGDDGQGVPAGG |
| C-terminal domain (CTD)<br>(residues 252 to 401) | ETGQVLAKSNDLDFNTMWVDPANSGIRRQLGDKALGGTVIYVNALGTHGLVVANSDQVNSNTWW<br>DAQDSITNPAHFDNEGKLYSDWRLPTRFELNLIYMMRNELGNFLAGNYWSSIEKSSANSWVFNS<br>KTGEIKDIAKSKTAAVRAVRAF |

### Supplementary Table 2 | X-ray data collection and refinement statistics

| | | |
|---|---|---|
| **Crystal Parameters** | | |
| Space group | *C*2 | *P*321 |
| Cell dimensions (Å; °) | *a*=91.60, *b*=52.87, *c*=97.27; α=γ=90, β=121.09 | *a*=74.91, *b*=74.91, *c*=94.97; α=β=90, γ=120 |
| **Data collection** | | |
| Beamline | DLS I03 | DLS I03 |
| Wavelength (Å) | 0.97969 | 0.97625 |
| Resolution (Å) | 45.56-1.90 (1.95-1.90) | 64.87-1.90 (1.93-1.90) |
| Unique observations | 34006 (2460) | 24856 (1235) |
| $R_{merge}$ (%) | 0.058 (0.148) | 0.074 (0.541) |
| $\langle I \rangle / \sigma I$ | 16.7 (6.7) | 28.7 (5.8) |
| Completeness (%) | 99.3 (97.3) | 100.0 (99.8) |
| Redundancy | 3.3 (2.8) | 19.3 (14.8) |
| Wilson B (Å) | 17.3 | 24.1 |
| **Phasing** | | |
| Figure of merit (accentric/centric) | 0.251/0.010 | - |
| Phasing power | 0.865 | - |
| **Refinement** | | |
| $R_{work}/R_{free}$ (%) | 15.8/18.8 | 16.0/18.7 |
| Number of protein residues | 387 | 262 |
| Number of water molecules | 397 | 194 |
| Number of ligands | 0 | 2 $SO_4^{2}$ |
| $\langle B \rangle$ protein | 19.1 | 27.8 |
| $\langle B \rangle$ waters | 28.3 | 35.8 |
| $\langle B \rangle$ ligands | - | 34.5 |
| rmsd stereochemistry | | |
| Bond lengths (Å) | 0.005 | 0.008 |
| Bond angles (°) | 1.180 | 1.473 |
| Ramachandran analysis | | |
| Residues in outlier regions (%) | 0.0 | 0.0 |
| Residues in favoured regions (%) | 97.7 | 98.8 |
| Residues in allowed regions (%) | 100 | 100 |

Numbers in parentheses refer to the outermost resolution shell.
$R_{sym} = \sum|I - \langle I \rangle|/\sum I$ where $I$ is the integrated intensity of a given reflection and $\langle I \rangle$ is the mean intensity of multiple corresponding symmetry-related reflections.
$R_{work} = \sum||F_o| - |F_c||/\sum F_o$ where $F_o$ and $F_c$ are the observed and calculated structure factors, respectively.
$R_{free} = R_{work}$ calculated using 10% random data excluded from the refinement.
rmsd stereochemistry is the deviation from ideal values.
Ramachandran analysis was carried out using Molprobity [8]

### Supplementary Table 3 | SAXS data and refinement statistics

| **SAXS data collection** | WT | R342A peak 1 | R342A peak 2 | E368A | K369A | K380A | K385A | D386A | K391A |
|---|---|---|---|---|---|---|---|---|---|
| Beamline | DLS B21 | DLS B21 | DLS B21 | DLS B21 | DLS B21 | DLS B21 | DLS B21 | DLS B21 | DLS B21 |
| Wavelength (Å) | 1.0 | 1.0 | 1.0 | 1.0 | 1.0 | 1.0 | 1.0 | 1.0 | 1.0 |
| q Range ($Å^{-1}$) | 0.003 to 0.44 | 0.003 to 0.44 | 0.003 to 0.44 | 0.003 to 0.44 | 0.003 to 0.44 | 0.003 to 0.44 | 0.003 to 0.44 | 0.003 to 0.44 | 0.003 to 0.44 |
| **Structural parameters** | | | | | | | | | |
| I(0) | 0.091 ± 5.6e-05 | 0.091 ± 2.3e-04 | 0.075 ± 4.5e-05 | 0.090 ± 5.4e-05 | 0.091 ± 5.2e-05 | 0.094 ± 8.2e-05 | 0.092 ± 1.0e-04 | 0.093 ± 6.3e-05 | 0.093 ± 5.2e-05 |
| $R_g$ (nm) (from Guinier) | 2.77 ± 0.10 | 2.68 ± 0.04 | 1.84 ± 0.15 | 2.79 ± 0.36 | 2.78 ± 0.18 | 2.79 ± 0.04 | 2.81 ± 0.18 | 2.83 ± 0.02 | 2.77 ± 0.01 |
| $R_g$ (nm) (from P(r)) | 2.85 ± 0.01 | 2.75 ± 0.01 | 1.85 ± 0.01 | 2.82 ± 0.01 | 2.85 ± 0.01 | 2.86 ± 0.01 | 2.88 ± 0.01 | 2.84 ± 0.01 | 2.83 ± 0.01 |
| $D_{max}$ (nm) (from P(r)) | 9.70 | 9.38 | 6.44 | 9.89 | 9.73 | 9.77 | 9.84 | 9.77 | 9.49 |
| MW (kDa) (from sequence) | 18549 | 18464 | 18464 | 18475 | 18492 | 18492 | 18492 | 18479 | 18492 |
| MW (kDa) (from SAXS) | 50877 | 47126 | 18809 | 50626 | 51962 | 51517 | 51667 | 51522 | 51493 |

### Supplementary Table 4 | SAXS ensemble optimization parameters.

| | |
|---|---|
| $R_{flex}$ (random) (%) | 88.10 (84.55) |
| $R_{sig}$ | 4.40 |
| $\chi^2$ | 1.05 |
| Cluster 1 (%) | 30 |
| Cluster 2 (%) | 20 |
| Cluster 3 (%) | 50 |
| Final ensemble $R_g$ (nm) | 2.82 |
| Final ensemble $D_{max}$ (nm) | 9.18 |

**Supplementary Table 5 | Lcl homologs from the DUF1566 gly_rich_SclB superfamily.**

| Description | Species | NIH accession code |
|---|---|---|
| DUF1566 domain-containing protein | Legionella pneumophila | WP_010948344.1 |
| DUF1566 domain-containing protein | Legionella pneumophila | WP_015961725.1 |
| DUF1566 domain-containing protein | Legionella pneumophila | WP_011216516.1 |
| Legionella vir region protein | Flavobacteria. bacterium BAL38 | EAZ94970.1 |
| Hypothetical protein GOS_6310005, partial (marine metagenome) | N/A | ECC71828.1 |
| DUF1566 domain-containing protein | Legionella pneumophila | WP_011947561.1 |
| DUF1566 domain-containing protein | Legionella pneumophila | WP_013101911.1 |
| DUF1566 domain-containing protein | Legionella pneumophila | WP_040185895.1 |
| DUF1566 domain-containing protein | Legionella pneumophila | WP_061563548.1 |
| DUF1566 domain-containing protein | Legionella pneumophila | WP_062725249.1 |
| HbP1 | Legionella pneumophila | ADN03210.1 |
| DUF1566 domain-containing protein | Legionella pneumophila | WP_014844754.1 |
| DUF1566 domain-containing protein | Legionella pneumophila | WP_015443987.1 |
| DUF1566 domain-containing protein | Legionella pneumophila | WP_016357032.1 |
| DUF1566 domain-containing protein, partial | Legionella pneumophila | WP_021459955.1 |
| DUF1566 domain-containing protein | Legionella oakridgensis | WP_025385719.1 |
| DUF1566 domain-containing protein | Legionella oakridgensis | WP_025385919.1 |
| DUF1566 domain-containing protein | Legionella oakridgensis | WP_035896162.1 |
| DUF1566 domain-containing protein, partial | Legionella pneumophila | WP_027264466.1 |
| DUF1566 domain-containing protein | Legionella pneumophila | WP_027267710.1 |
| DUF1566 domain-containing protein, partial | Legionella pneumophila | WP_032828702.1 |
| DUF1566 domain-containing protein, partial | Legionella pneumophila | WP_032828931.1 |
| DUF1566 domain-containing protein, partial | Legionella pneumophila | WP_032832228.1 |
| DUF1566 domain-containing protein, partial | Legionella pneumophila | WP_032833288.1 |
| DUF1566 domain-containing protein, partial | Legionella pneumophila | WP_038839654.1 |
| DUF1566 domain-containing protein, partial | Legionella pneumophila | WP_040146561.1 |
| DUF1566 domain-containing protein, partial | Legionella pneumophila | WP_040147717.1 |
| DUF1566 domain-containing protein | Legionella pneumophila | WP_040164635.1 |
| DUF1566 domain-containing protein, partial | Legionella pneumophila | WP_040176715.1 |
| DUF1566 domain-containing protein, partial | Legionella pneumophila | WP_040183632.1 |
| DUF1566 domain-containing protein | Legionella pneumophila | WP_040184835.1 |
| DUF1566 domain-containing protein, partial | Legionella pneumophila | WP_040187765.1 |
| DUF1566 domain-containing protein, partial | Legionella pneumophila | WP_040190847.1 |
| DUF1566 domain-containing protein | Legionella pneumophila | WP_042237472.1 |
| DUF1566 domain-containing protein, partial | Legionella pneumophila | WP_042637136.1 |
| DUF1566 domain-containing protein, partial | Legionella pneumophila | WP_042645969.1 |
| DUF1566 domain-containing protein | Legionella pneumophila | WP_042647776.1 |
| DUF1566 domain-containing protein, partial | Legionella pneumophila | WP_042647794.1 |
| DUF1566 domain-containing protein, partial | Legionella pneumophila | WP_042647975.1 |
| DUF1566 domain-containing protein, partial | Legionella pneumophila | WP_042650802.1 |
| DUF1566 domain-containing protein, partial | Legionella pneumophila | WP_042655488.1 |
| DUF1566 domain-containing protein | Legionella pneumophila | WP_042754707.1 |
| DUF1566 domain-containing protein | Legionella pneumophila | WP_059428048.1 |
| DUF1566 domain-containing protein | Legionella pneumophila | WP_059428638.1 |
| DUF1566 domain-containing protein, partial | Legionella pneumophila | WP_044499023.1 |
| DUF1566 domain-containing protein, partial | Legionella pneumophila | WP_044552931.1 |
| DUF1566 domain-containing protein, partial | Legionella pneumophila | WP_047661805.1 |
| DUF1566 domain-containing protein, partial | Legionella pneumophila | WP_059411200.1 |
| DUF1566 domain-containing protein, partial | Legionella pneumophila | WP_059431367.1 |
| DUF1566 domain-containing protein | Legionella pneumophila | WP_060871594.1 |
| DUF1566 domain-containing protein | Legionella pneumophila | WP_061469204.1 |
| DUF1566 domain-containing protein | Legionella pneumophila | WP_061684452.1 |
| DUF1566 domain-containing protein | Legionella pneumophila | WP_061691911.1 |
| DUF1566 domain-containing protein | Legionella pneumophila | WP_061635485.1 |
| DUF1566 domain-containing protein | Legionella pneumophila | WP_061710309.1 |
| DUF1566 domain-containing protein | Legionella pneumophila | WP_061638011.1 |
| DUF1566 domain-containing protein | Legionella pneumophila | WP_061832330.1 |
| DUF1566 domain-containing protein | Legionella pneumophila | WP_061515237.1 |

| | | |
|---|---|---|
| DUF1566 domain-containing protein | Legionella pneumophila | WP_230305795.1 |
| DUF1566 domain-containing protein | Legionella pneumophila | WP_061690779.1 |
| DUF1566 domain-containing protein | Legionella pneumophila | WP_061722744.1 |
| DUF1566 domain-containing protein | Legionella pneumophila | WP_061722072.1 |
| DUF1566 domain-containing protein | Legionella pneumophila | WP_061637081.1 |
| DUF1566 domain-containing protein | Legionella pneumophila | WP_061723125.1 |
| DUF1566 domain-containing protein | Legionella pneumophila | WP_061716791.1 |
| DUF1566 domain-containing protein | Legionella pneumophila | WP_061836912.1 |
| DUF1566 domain-containing protein | Legionella pneumophila | WP_061639199.1 |
| DUF1566 domain-containing protein | Legionella pneumophila | WP_061723636.1 |
| DUF1566 domain-containing protein | Legionella pneumophila | WP_061398413.1 |
| DUF1566 domain-containing protein | Legionella pneumophila | WP_061684485.1 |
| DUF1566 domain-containing protein | Legionella pneumophila | WP_061828234.1 |
| DUF1566 domain-containing protein | Legionella pneumophila | WP_061635538.1 |
| DUF1566 domain-containing protein | Legionella pneumophila | WP_061723696.1 |
| DUF1566 domain-containing protein | Legionella pneumophila | WP_061828204.1 |
| DUF1566 domain-containing protein | Legionella pneumophila | WP_061827839.1 |
| DUF1566 domain-containing protein | Legionella pneumophila | WP_061828230.1 |
| DUF1566 domain-containing protein | Legionella pneumophila | WP_061724091.1 |
| DUF1566 domain-containing protein | Legionella pneumophila | WP_061467335.1 |
| DUF1566 domain-containing protein | Legionella pneumophila | WP_061773537.1 |
| DUF1566 domain-containing protein | Legionella pneumophila | WP_061467096.1 |
| DUF1566 domain-containing protein | Legionella pneumophila | WP_061480980.1 |
| DUF1566 domain-containing protein | Legionella pneumophila | WP_061481517.1 |
| DUF1566 domain-containing protein | Legionella pneumophila | WP_061483720.1 |
| DUF1566 domain-containing protein | Legionella pneumophila | WP_061512424.1 |
| DUF1566 domain-containing protein | Legionella pneumophila | WP_061513719.1 |
| DUF1566 domain-containing protein | Legionella pneumophila | WP_061773165.1 |
| DUF1566 domain-containing protein | Legionella pneumophila | WP_061784476.1 |
| DUF1566 domain-containing protein, partial | Legionella pneumophila | WP_062721381.1 |
| DUF1566 domain-containing protein, partial | Legionella pneumophila | WP_062739669.1 |
| DUF1566 domain-containing protein, partial | Legionella pneumophila | WP_062739721.1 |
| DUF1566 domain-containing protein, partial | Legionella pneumophila | WP_062756802.1 |
| DUF1566 domain-containing protein, partial | Legionella pneumophila | WP_062777099.1 |
| DUF1566 domain-containing protein | Legionella pneumophila | WP_062896185.1 |
| DUF1566 domain-containing protein | Legionella oakridgensis | WP_064101346.1 |
| DUF1566 domain-containing protein | Legionella pneumophila | WP_070322485.1 |
| DUF1566 domain-containing protein | Legionella pneumophila | WP_070322308.1 |
| MAG: hypothetical protein A3D92_17110 | Bacteroidetes bacterium RIFCSPHIGHO2_02_FULL_44_7 | OFZ16950.1 |
| MAG: hypothetical protein A3D31_13075 | Candidatus Fluviicola riflensis | OGS77914.1 |
| DUF1566 domain-containing protein, partial | Legionella pneumophila | WP_143460085.1 |
| DUF1566 domain-containing protein, partial | Legionella pneumophila | WP_143460548.1 |
| MAG: hypothetical protein COA38_21500 | Fluviicola sp. | PHR16987.1 |
| collagen triple helix repeat protein | Comamonas sp. 26 | PIG08876.1 |
| MAG: hypothetical protein CK423_04720 | Legionella sp. | PJD94984.1 |
| MAG: hypothetical protein CK426_01620 | Legionella sp. | PJD99773.1 |
| DUF1566 domain-containing protein | Nonlabens sp. MB-3u-79 | WP_101013415.1 |
| DUF1566 domain-containing protein | Legionella pneumophila | WP_106174748.1 |
| DUF1566 domain-containing protein | Legionella pneumophila | WP_106184284.1 |
| DUF1566 domain-containing protein | Legionella pneumophila | WP_106195271.1 |
| DUF1566 domain-containing protein | Legionella pneumophila | WP_106225763.1 |
| DUF1566 domain-containing protein | Legionella pneumophila | WP_106229744.1 |
| DUF1566 domain-containing protein | Legionella pneumophila | WP_110570296.1 |
| DUF1566 domain-containing protein | Oceanihabitans sediminis | WP_123796521.1 |
| DUF1566 domain-containing protein | Oceanihabitans sediminis | RCU56654.1 |
| DUF1566 domain-containing protein | Legionella pneumophila | WP_114693300.1 |
| DUF1566 domain-containing protein, partial | Legionella pneumophila | WP_114674093.1 |
| tail fiber protein | Legionella longbeachae | STY20446.1 |
| DUF1566 domain-containing protein | Legionella quateirensis | WP_115148663.1 |
| uncharacterized protein DUF1566, partial | Winogradskyella pacifica | REE25412.1 |
| DUF1566 domain-containing protein | Legionella pneumophila | WP_120061973.1 |
| DUF1566 domain-containing protein | Legionella pneumophila | WP_134984243.1 |
| DUF1566 domain-containing protein | Legionella pneumophila | WP_129368547.1 |

| | | |
|---|---|---|
| DUF1566 domain-containing protein | Legionella quateirensis | WP_133141067.1 |
| DUF1566 domain-containing protein, partial | Legionella pneumophila | WP_134921750.1 |
| DUF1566 domain-containing protein | Legionella pneumophila | WP_136669451.1 |
| DUF1566 domain-containing protein, partial | Legionella pneumophila | WP_136691115.1 |
| DUF1566 domain-containing protein | Legionella pneumophila | WP_136627821.1 |
| DUF1566 domain-containing protein | Legionella pneumophila | WP_136633845.1 |
| DUF1566 domain-containing protein | Legionella pneumophila | WP_136648505.1 |
| MAG: DUF1566 domain-containing protein, partial | Gammaproteobacteria bacterium | TLZ59698.1 |
| MAG: DUF1566 domain-containing protein | Chitinophagaceae bacterium | TVR83543.1 |
| DUF1566 domain-containing protein, partial | Legionella pneumophila | WP_148654818.1 |
| hypothetical protein EZS27_000721<br>(termite gut metagenome) | N/A | KAA6351924.1 |
| Hypothetical protein | Cupriavidus sp. SK-3 | WP_152561645.1 |
| DUF1566 domain-containing protein, partial | Legionella pneumophila | WP_154231773.1 |
| unannotated protein (freshwater metagenome) | N/A | CAB4331133.1 |
| DUF1566 domain-containing protein, partial | Legionella pneumophila | WP_154694957.1 |
| DUF1566 domain-containing protein, partial | Legionella pneumophila | WP_154698949.1 |
| DUF1566 domain-containing protein, partial | Legionella pneumophila | WP_154698961.1 |
| DUF1566 domain-containing protein, partial | Legionella pneumophila | WP_154699083.1 |
| DUF1566 domain-containing protein, partial | Legionella pneumophila | WP_154699070.1 |
| DUF1566 domain-containing protein, partial | Legionella pneumophila | WP_154699188.1 |
| DUF1566 domain-containing protein, partial | Legionella pneumophila | WP_154699211.1 |
| DUF1566 domain-containing protein, partial | Legionella pneumophila | WP_154699232.1 |
| DUF1566 domain-containing protein, partial | Legionella pneumophila | WP_154699253.1 |
| DUF1566 domain-containing protein, partial | Legionella pneumophila | WP_154699450.1 |
| DUF1566 domain-containing protein, partial | Legionella pneumophila | WP_154699477.1 |
| DUF1566 domain-containing protein, partial | Legionella pneumophila | WP_154699519.1 |
| DUF1566 domain-containing protein, partial | Legionella pneumophila | WP_154699508.1 |
| DUF1566 domain-containing protein, partial | Legionella pneumophila | WP_154699564.1 |
| DUF1566 domain-containing protein, partial | Legionella pneumophila | WP_154699554.1 |
| DUF1566 domain-containing protein, partial | Legionella pneumophila | WP_154699579.1 |
| DUF1566 domain-containing protein, partial | Legionella pneumophila | WP_154699623.1 |
| DUF1566 domain-containing protein, partial | Legionella pneumophila | WP_154699798.1 |
| DUF1566 domain-containing protein, partial | Legionella pneumophila | WP_161503312.1 |
| DUF1566 domain-containing protein, partial | Legionella pneumophila | WP_161503617.1 |
| DUF1566 domain-containing protein, partial | Legionella pneumophila | WP_161505509.1 |
| DUF1566 domain-containing protein, partial | Legionella pneumophila | WP_161513085.1 |
| DUF1566 domain-containing protein, partial | Legionella pneumophila | WP_161513358.1 |
| DUF1566 domain-containing protein, partial | Legionella pneumophila | WP_161513925.1 |
| DUF1566 domain-containing protein | Legionella pneumophila | WP_161514298.1 |
| DUF1566 domain-containing protein, partial | Legionella pneumophila | WP_161514748.1 |
| DUF1566 domain-containing protein, partial | Legionella pneumophila | WP_161516834.1 |
| Collagen triple helix repeat<br>(uncultured Caudovirales phage) | N/A | CAB4171944.1 |
| DUF1566 domain-containing protein | Winogradskyella wichelsiae | WP_179375955.1 |
| hypothetical protein GCM10010832_23760 | Psychroflexus planctonicus | GGE43047.1 |
| DUF1566 domain-containing protein | Psychroflexus planctonicus | WP_194428404.1 |
| phage tail protein | Legionella pneumophila | WP_281162213.1 |
| DUF1566 domain-containing protein, partial | Legionella pneumophila | WP_247337804.1 |
| DUF1566 domain-containing protein, partial | Legionella pneumophila | WP_247386728.1 |
| DUF1566 domain-containing protein | Legionella pneumophila | WP_241505424.1 |
| DUF1566 domain-containing protein | Legionella pneumophila | WP_241507376.1 |
| DUF1566 domain-containing protein, partial | Legionella pneumophila | WP_247358519.1 |
| hypothetical protein | Comamonas sp. 26 | WP_199173708.1 |
| DUF1566 domain-containing protein | Legionella sp. PATHC032 | WP_265719455.1 |
| DUF1566 domain-containing protein, partial | Legionella pneumophila | WP_247094671.1 |
| DUF1566 domain-containing protein, partial | Legionella pneumophila | WP_241510043.1 |
| DUF1566 domain-containing protein, partial | Legionella pneumophila | WP_241508403.1 |
| DUF1566 domain-containing protein | Methylomonas paludis | WP_215584844.1 |
| DUF1566 domain-containing protein, partial | Legionella pneumophila | WP_229309814.1 |
| DUF1566 domain-containing protein, partial | Legionella pneumophila | WP_230308248.1 |
| DUF1566 domain-containing protein, partial | Legionella pneumophila | WP_230314272.1 |
| DUF1566 domain-containing protein | Legionella pneumophila | WP_236715301.1 |
| hypothetical protein | Legionella bononiensis | WP_238400292.1 |

| | | |
|---|---|---|
| DUF1566 domain-containing protein | Legionella bononiensis | WP_238400409.1 |
| DUF1566 domain-containing protein, partial | Legionella quateirensis | WP_238585567.1 |
| DUF1566 domain-containing protein | Legionella pneumophila | WP_241507749.1 |
| DUF1566 domain-containing protein, partial | Legionella pneumophila | WP_241534195.1 |
| DUF1566 domain-containing protein, partial | Legionella pneumophila | WP_241506022.1 |
| DUF1566 domain-containing protein, partial | Legionella pneumophila | WP_241505578.1 |
| DUF1566 domain-containing protein | Legionella pneumophila | WP_241533060.1 |
| DUF1566 domain-containing protein | Legionella pneumophila | WP_241535769.1 |
| | Legionella pneumophila | WP_241534112.1 |
| DUF1566 domain-containing protein, partial | Legionella pneumophila | WP_241534982.1 |
| DUF1566 domain-containing protein | Legionella pneumophila | WP_241532330.1 |
| DUF1566 domain-containing protein | Thiocystis minor | WP_242523155.1 |
| hypothetical protein | Methylobacter sp. S3L5C | WP_243219470.1 |
| DUF1566 domain-containing protein, partial | Legionella pneumophila | WP_247356958.1 |
| phage tail protein | Legionella pneumophila | WP_269568483.1 |
| DUF1566 domain-containing protein, partial | Legionella pneumophila | WP_269562955.1 |
| hypothetical protein | Winogradskyella pacifica | WP_257018464.1 |
| hypothetical protein | Winogradskyella wichelsiae | WP_262886520.1 |
| hypothetical protein | Bacteroidia bacterium | GHS95945.1 |
| phage tail protein | Legionella pneumophila | WP_265665447.1 |
| phage tail protein | Legionella sp. PATHC039 | WP_265732188.1 |
| phage tail protein | Legionella pneumophila | WP_265672815.1 |
| phage tail protein | Legionella pneumophila | WP_265672181.1 |
| phage tail protein | Legionella pneumophila | WP_269568141.1 |
| phage tail protein, partial | Legionella pneumophila | WP_269563856.1 |
| DUF1566 domain-containing protein, partial | Legionella pneumophila | WP_269568104.1 |
| phage tail protein | Legionella pneumophila | WP_269568937.1 |
| DUF1566 domain-containing protein, partial | Legionella pneumophila | WP_269567031.1 |
| phage tail protein | Legionella pneumophila | WP_269564737.1 |
| phage tail protein | Legionella pneumophila | WP_269564955.1 |
| DUF1566 domain-containing protein, partial | Legionella pneumophila | WP_269565042.1 |
| phage tail protein, partial | Legionella pneumophila | WP_269566652.1 |
| DUF1566 domain-containing protein, partial | Legionella pneumophila | WP_269566086.1 |
| DUF1566 domain-containing protein, partial | Legionella pneumophila | WP_269569356.1 |
| DUF1566 domain-containing protein, partial | Legionella pneumophila | WP_269562660.1 |
| DUF1566 domain-containing protein, partial | Legionella pneumophila | WP_269568473.1 |
| phage tail protein | Legionella pneumophila | WP_269568720.1 |
| DUF1566 domain-containing protein, partial | Legionella pneumophila | WP_272838090.1 |
| DUF1566 domain-containing protein, partial | Legionella pneumophila | WP_272850246.1 |
| MAG: Uncharacterised protein | Formosa sp. Hel1_33_131 | CAI8189797.1 |
| phage tail protein | Legionella pneumophila | WP_281489849.1 |
| phage tail protein | Legionella pneumophila | WP_283324708.1 |

**Supplementary Table 6 | Average frequency of occurrence of Lcl-CTD/C4S contacts during MD simulations.**

| **Residue**[a] | **Chain A**[b] **(%)** | **Chain B**[b] **(%)** | **Chain C**[b] **(%)** |
|---|---|---|---|
| S371 | 40.9 | 40.3 | 37.6 |
| A372 | 29.2 | 31.1 | 25.7 |
| **K391** | 32.4 | 24.8 | 19.3 |
| S390 | 20.9 | 36.5 | 17.2 |
| A388 | 21.9 | 22.6 | 18.2 |
| S370 | 26.9 | 14.7 | 20.7 |
| N373 | 24.4 | 11.6 | 14.5 |
| W315 | 11.1 | 22.3 | 9.6 |
| I387 | 14.8 | 4.2 | 7.1 |
| **K385** | 16.3 | 1.8 | 4.8 |
| **K369** | 13.4 | 3.1 | 3.0 |
| T392 | 10.1 | 0.8 | 7.5 |
| N362 | 13.4 | 0.4 | 3.7 |
| N378 | 10.6 | 0.3 | 0.3 |
| **K380** | 10.2 | 0.2 | 0.1 |

[a]Residues are sorted by decreasing average frequency. Lysines are highlighted in bold.
[b]The frequency was calculated as average over all the replicas (production only). Only residues with a frequency ≥ 10% for at least one chain are reported.

**Supplementary Table 7 | Average frequency of occurrence of Lcl-CTD/C4S hydrogen bonds during MD simulations.**

| Residue[a] | Chain A[b] (%) | Chain B[b] (%) | Chain C[b] (%) |
|---|---|---|---|
| S371 | 39.7 | 36.8 | 40.8 |
| S390 | 11.7 | 27.8 | 10.1 |
| **K391** | 19.1 | 11.7 | 9.7 |
| T392 | 15.9 | 1.9 | 11.8 |
| N373 | 10.6 | 7.8 | 8.4 |
| S370 | 12.0 | 4.2 | 4.9 |
| N362 | 14.6 | 0.7 | 4.6 |
| **K385** | 11.8 | 0.6 | 2.9 |

[a]Residues are sorted by decreasing average frequency. Lysines are highlighted in bold. Hydrogen bonding interactions were detected using Visual Molecular Dynamics[9] with a Donor-Acceptor distance threshold of 3.5 Å and a Hydrogen-Donor-Acceptor angle of 30°.
[b]The frequency was calculated as average over all the replicas (production only). When more than one donor/acceptor per residue was involved in hydrogen bonding, the frequency values were added together. Only residues with a frequency >= 10% for at least one chain are reported.

### Supplementary Table 8 | Primers, plasmids and strains used in this study

| Primer | Description | Sequence (5' to 3') |
|---|---|---|
| *lcl*-UpF | Gene KO – *L. pneumophila* 130b *lcl* upstream F | ATACGTACACGAAGGCGACC |
| *lcl* -UpR | Gene KO – *L. pneumophila* 130b *lcl* upstream R | TCTTTATTTCCCACTATATGCTCACTTG |
| *lcl*-DownF | Gene KO – *L. pneumophila* 130b *lcl* downstream F | GTATGGCAAAACTTCAATTTTGCT |
| *lcl* -DownR | Gene KO – *L. pneumophila* 130b *lcl* downstream R | AGAAGTACGCCCTGGTTTCG |
| *lcl*-P1 | Gene KO - Kn-resistance cassette F | AAATAAAAGAGACAAGTGAGCATATAGTGG GAAATAAAGAGTGTAGGCTGGAGCTGCTTC |
| *lcl*-P2 | Gene KO - Kn-resistance cassette R | GAACATTTTCATCCTTGAGCAAAATTGAAGT TTTGCCATACCATATGAATATCCTCCTTA |
| RLC1 | Recombinant expression of *L. pneumophila* 130b Lcl F | GACGACGACAAGATGAAAAGCAATCCGGCC TCGC |
| RLC2 | Recombinant expression of *L. pneumophila* 130b Lcl R | GAGGAGAAGCCCGGttaAAAGGCTCTTACAGC ACGTAC |
| RLC3 | Recombinant expression of *L. pneumophila* 130b Lcl-CTD F | GACGACGACAAGATGGAAACGGGACAAGTC CTTGC |
| RLC4 | Recombinant expression of *L. pneumophila* 130b Lcl-CTD R | GAGGAGAAGCCCGGttaAAAGGCTCTTACAGC ACGTAC |

| Plasmid | Description | Reference |
|---|---|---|
| pET46 Ek/LIC | Expression vector | Novagen |
| pRLC1 | pET46 Ek/LIC expression of *L. pneumophila* Lcl residues 1-401 | This study |
| pRLC2 | pET46 Ek/LIC expression of *L. pneumophila* Lcl CTD residues 252-401 | This study |
| pET28b | Expression vector | Novagen |
| pRLCm1 | pET28b expression of *L. pneumophila* Lcl (CTD) residues 252-401 containing R342A mutation | This study |
| pRLCm2 | pET28b expression of *L. pneumophila* Lcl (CTD)residues 252-401 containing E368A mutation | This study |
| pRLCm3 | pET28b expression of *L. pneumophila* Lcl (CTD)residues 252-401 containing K369A mutation | This study |
| pRLCm4 | pET28b expression of *L. pneumophila* Lcl (CTD)residues 252-401 containing K380A mutation | This study |
| pRLCm5 | pET28b expression of *L. pneumophila* Lcl (CTD)residues 252-401 containing K385A mutation | This study |
| pRLCm6 | pET28b expression of *L. pneumophila* Lcl (CTD)residues 252-401 containing D386A mutation | This study |
| pRLCm7 | pET28b expression of *L. pneumophila* Lcl (CTD)residues 252-401 containing K391A mutation | This study |

| Bacterial strain | Serogroup/ genotype | Reference |
|---|---|---|
| *L. pneumophila* 130b | ATCC strain BAA-74. Served as the wild type and parent for all mutants. | [10,11] |
| *L. pneumophila* NU468 | *130b strain carrying lcl mutation* | This study |
| *L. pneumophila* NU469 | *130b strain carrying lcl mutation* | This study |
| *L. pneumophila* NU275 | *130b strain carrying lspF (T2SS) mutation* | [12] |
| *E. coli* DH5a | *F- Φ80lacZΔM15 Δ(lacZYA-argF) U169 recA1 endA1 hsdR17(rk-, mk+) phoA supE44 thi-1 gyrA96 relA1 λ-* | Invitrogen |

| | | |
|---|---|---|
| *E. coli* BL21 (DE3) | *fhuA2 [lon] ompT gal (λ DE3) [dcm] ΔhsdS*<br>*λ DE3 = λ sBamHIo ΔEcoRI-B int::(lacI::PlacUV5::T7 gene1) i21 Δnin5* | NEB |

### Supplementary Table 9 | Synthetic genes

| Gene | Description | Sequence (5' to 3') |
|---|---|---|
| gRLCm1 | Lcl-CTD R342A mutant | CCATGGTCCATCATCATCATCATCATGTGGACGACGACGACAAGATGGAAACGGGACAA<br>GTCCTTGCCAAATCAAATGACCTCGATTTCAATACCATGTGGGTTGACCCGGCAAATTC<br>CGGCATTAGGCGGCAATTAGGCGATAAAGCTCTTGGCGGTACAGTGATTTATGTTAATG<br>CGCTAGGTACACACGGGCTTGTAGTAGCAAATTCAGATCAAGTTAATTCAAACACATGG<br>TGGGATGCTCAAGATTCCATAACAAATCCTGCCCACTTTGACAATGAAGGTAAATTATA<br>TTCTGACTGGAGGCTCCCTACCGCGTTTGAATTGAATTTAATTTATATGATGCGAAATG<br>AGCTGGGAAATTTTTTGGCAGGTAATTACTGGAGTTCGATTGAAAAATCATCGGCAAAC<br>AGCTGGGTATTTAATTCTAAAACAGGGGAAATTAAAGACATTGCCAAAAGTAAAACGGC<br>TGCTGTACGTGCTGTAAGAGCCTTTTAACTCGAG |
| gRLCm2 | Lcl-CTD E368A mutant | CCATGGTCCATCATCATCATCATCATGTGGACGACGACGACAAGATGGAAACGGGACAA<br>GTCCTTGCCAAATCAAATGACCTCGATTTCAATACCATGTGGGTTGACCCGGCAAATTC<br>CGGCATTAGGCGGCAATTAGGCGATAAAGCTCTTGGCGGTACAGTGATTTATGTTAATG<br>CGCTAGGTACACACGGGCTTGTAGTAGCAAATTCAGATCAAGTTAATTCAAACACATGG<br>TGGGATGCTCAAGATTCCATAACAAATCCTGCCCACTTTGACAATGAAGGTAAATTATA<br>TTCTGACTGGAGGCTCCCTACCCGTTTTGAATTGAATTTAATTTATATGATGCGAAATG<br>AGCTGGGAAATTTTTTGGCAGGTAATTACTGGAGTTCGATTGCGAAATCATCGGCAAAC<br>AGCTGGGTATTTAATTCTAAAACAGGGGAAATTAAAGACATTGCCAAAAGTAAAACGGC<br>TGCTGTACGTGCTGTAAGAGCCTTTTAACTCGAG |
| gRLCm3 | Lcl-CTD K369A mutant | CCATGGTCCATCATCATCATCATCATGTGGACGACGACGACAAGATGGAAACGGGACAA<br>GTCCTTGCCAAATCAAATGACCTCGATTTCAATACCATGTGGGTTGACCCGGCAAATTC<br>CGGCATTAGGCGGCAATTAGGCGATAAAGCTCTTGGCGGTACAGTGATTTATGTTAATG<br>CGCTAGGTACACACGGGCTTGTAGTAGCAAATTCAGATCAAGTTAATTCAAACACATGG<br>TGGGATGCTCAAGATTCCATAACAAATCCTGCCCACTTTGACAATGAAGGTAAATTATA<br>TTCTGACTGGAGGCTCCCTACCCGTTTTGAATTGAATTTAATTTATATGATGCGAAATG<br>AGCTGGGAAATTTTTTGGCAGGTAATTACTGGAGTTCGATTGAAGCGTCATCGGCAAAC<br>AGCTGGGTATTTAATTCTAAAACAGGGGAAATTAAAGACATTGCCAAAAGTAAAACGGC<br>TGCTGTACGTGCTGTAAGAGCCTTTTAACTCGAG |
| gRLCm4 | Lcl-CTD K380A mutant | CCATGGTCCATCATCATCATCATCATGTGGACGACGACGACAAGATGGAAACGGGACAA<br>GTCCTTGCCAAATCAAATGACCTCGATTTCAATACCATGTGGGTTGACCCGGCAAATTC<br>CGGCATTAGGCGGCAATTAGGCGATAAAGCTCTTGGCGGTACAGTGATTTATGTTAATG<br>CGCTAGGTACACACGGGCTTGTAGTAGCAAATTCAGATCAAGTTAATTCAAACACATGG<br>TGGGATGCTCAAGATTCCATAACAAATCCTGCCCACTTTGACAATGAAGGTAAATTATA<br>TTCTGACTGGAGGCTCCCTACCCGTTTTGAATTGAATTTAATTTATATGATGCGAAATG<br>AGCTGGGAAATTTTTTGGCAGGTAATTACTGGAGTTCGATTGAAAAATCATCGGCAAAC<br>AGCTGGGTATTTAATTCTGCGACAGGGGAAATTAAAGACATTGCCAAAAGTAAAACGGC<br>TGCTGTACGTGCTGTAAGAGCCTTTTAACTCGAG |
| gRLCm5 | Lcl-CTD K385A mutant | CCATGGTCCATCATCATCATCATCATGTGGACGACGACGACAAGATGGAAACGGGACAA<br>GTCCTTGCCAAATCAAATGACCTCGATTTCAATACCATGTGGGTTGACCCGGCAAATTC<br>CGGCATTAGGCGGCAATTAGGCGATAAAGCTCTTGGCGGTACAGTGATTTATGTTAATG<br>CGCTAGGTACACACGGGCTTGTAGTAGCAAATTCAGATCAAGTTAATTCAAACACATGG<br>TGGGATGCTCAAGATTCCATAACAAATCCTGCCCACTTTGACAATGAAGGTAAATTATA<br>TTCTGACTGGAGGCTCCCTACCCGTTTTGAATTGAATTTAATTTATATGATGCGAAATG<br>AGCTGGGAAATTTTTTGGCAGGTAATTACTGGAGTTCGATTGAAAAATCATCGGCAAAC<br>AGCTGGGTATTTAATTCTAAAACAGGGGAAATTGCGGACATTGCCAAAAGTAAAACGGC<br>TGCTGTACGTGCTGTAAGAGCCTTTTAACTCGAG |
| gRLCm6 | Lcl-CTD D386A mutant | CCATGGTCCATCATCATCATCATCATGTGGACGACGACGACAAGATGGAAACGGGACAA<br>GTCCTTGCCAAATCAAATGACCTCGATTTCAATACCATGTGGGTTGACCCGGCAAATTC<br>CGGCATTAGGCGGCAATTAGGCGATAAAGCTCTTGGCGGTACAGTGATTTATGTTAATG<br>CGCTAGGTACACACGGGCTTGTAGTAGCAAATTCAGATCAAGTTAATTCAAACACATGG<br>TGGGATGCTCAAGATTCCATAACAAATCCTGCCCACTTTGACAATGAAGGTAAATTATA<br>TTCTGACTGGAGGCTCCCTACCCGTTTTGAATTGAATTTAATTTATATGATGCGAAATG<br>AGCTGGGAAATTTTTTGGCAGGTAATTACTGGAGTTCGATTGAAAAATCATCGGCAAAC |

| | | |
|---|---|---|
| | | AGCTGGGTATTTAATTCTAAAACAGGGGAAATTAAAGCGATTGCCAAAAGTAAAACGGC<br>TGCTGTACGTGCTGTAAGAGCCTTTTAACTCGAG |
| gRLCm7 | Lcl-CTD K391A mutant | CCATGGTCCATCATCATCATCATCATGTGGACGACGACGACAAGATGGAAACGGGACAA<br>GTCCTTGCCAAATCAAATGACCTCGATTTCAATACCATGTGGGTTGACCCGGCAAATTC<br>CGGCATTAGGCGGCAATTAGGCGATAAAGCTCTTGGCGGTACAGTGATTTATGTTAATG<br>CGCTAGGTACACACGGGCTTGTAGTAGCAAATTCAGATCAAGTTAATTCAAACACATGG<br>TGGGATGCTCAAGATTCCATAACAAATCCTGCCCACTTTGACAATGAAGGTAAATTATA<br>TTCTGACTGGAGGCTCCCTACCCGTTTTGAATTGAATTTAATTTATATGATGCGAAATG<br>AGCTGGGAAATTTTTTGGCAGGTAATTACTGGAGTTCGATTGAAAAATCATCGGCAAAC<br>AGCTGGGTATTTAATTCTAAAACAGGGGAAATTAAAGACATTGCCAAAAGTGCGACGGC<br>TGCTGTACGTGCTGTAAGAGCCTTTTAACTCGAG |

*Restriction sites are underlined

### References


1 Camilloni, C., De Simone, A., Vranken, W. F. & Vendruscolo, M. Determination of secondary structure populations in disordered states of proteins using nuclear magnetic resonance chemical shifts. *Biochemistry* **51**, 2224-2231, doi:10.1021/bi3001825 (2012).

2 Holm, L. & Sander, C. Dali: a network tool for protein structure comparison. *Trends Biochem Sci* **20**, 478-480, doi:10.1016/s0968-0004(00)89105-7 (1995).

3 Huang, K. F. *et al.* Crystal structure of a platelet-agglutinating factor isolated from the venom of Taiwan habu (Trimeresurus mucrosquamatus). *Biochem J* **378**, 399-407, doi:10.1042/BJ20031507 (2004).

4 Horii, K., Okuda, D., Morita, T. & Mizuno, H. Crystal structure of EMS16 in complex with the integrin alpha2-I domain. *J Mol Biol* **341**, 519-527, doi:10.1016/j.jmb.2004.06.036 (2004).

5 Luo, Y. *et al.* Crystal structure of enteropathogenic Escherichia coli intimin-receptor complex. *Nature* **405**, 1073-1077, doi:10.1038/35016618 (2000).

6 Hamburger, Z. A., Brown, M. S., Isberg, R. R. & Bjorkman, P. J. Crystal structure of invasin: a bacterial integrin-binding protein. *Science* **286**, 291-295 (1999).

7 Gautier, R., Douguet, D., Antonny, B. & Drin, G. HELIQUEST: a web server to screen sequences with specific alpha-helical properties. *Bioinformatics* **24**, 2101-2102, doi:10.1093/bioinformatics/btn392 (2008).

8 Davis, I. W., Murray, L. W., Richardson, J. S. & Richardson, D. C. MOLPROBITY: structure validation and all-atom contact analysis for nucleic acids and their complexes. *Nucleic Acids Res* **32**, W615-619, doi:10.1093/nar/gkh398 (2004).

9 Humphrey, W., Dalke, A. & Schulten, K. VMD: visual molecular dynamics. *J Mol Graph* **14**, 33-38, 27-38, doi:10.1016/0263-7855(96)00018-5 (1996).

10 DebRoy, S., Dao, J., Soderberg, M., Rossier, O. & Cianciotto, N. P. Legionella pneumophila type II secretome reveals unique exoproteins and a chitinase that promotes bacterial persistence in the lung. *Proc Natl Acad Sci U S A* **103**, 19146-19151, doi:10.1073/pnas.0608279103 (2006).

11 Schroeder, G. N. *et al.* Legionella pneumophila strain 130b possesses a unique combination of type IV secretion systems and novel Dot/Icm secretion system effector proteins. *J Bacteriol* **192**, 6001-6016, doi:10.1128/JB.00778-10 (2010).

12 Rossier, O., Starkenburg, S. R. & Cianciotto, N. P. Legionella pneumophila type II protein secretion promotes virulence in the A/J mouse model of Legionnaires' disease pneumonia. *Infect Immun* **72**, 310-321, doi:10.1128/IAI.72.1.310-321.2004 (2004).
